# Dual role of Human antigen R in Dengue virus infection: suppression of replication and promotion of cap-independent translation

**DOI:** 10.1101/2025.07.24.666571

**Authors:** Ashish Aneja, Risabh Sahu, Srishti Rajkumar Mishra, Arvind M. Korwar, E Sreekumar, Saumitra Das

**Author notes:** **Corresponding author- Saumitra Das**.

## Abstract

Host RNA-binding proteins (RBPs) play a pivotal role in regulating dengue virus (DENV) translation and replication through interactions with untranslated regions (UTRs) of viral RNA. We investigated host proteins associated with detergent-resistant membranes (DRMs) of the DENV replication complex and identified Human antigen R (HuR) as a key RBP enriched in the DRM. HuR was found to negatively regulate DENV replication by binding the DENV-3′UTR and impeding the association of polypyrimidine tract-binding protein (PTB), a known RNA stabilizer. Additionally, infection-induced modulation of HuR stabilized host mRNAs involved in innate immunity. Interestingly, *in vivo* validation in AG129 mice model highlights an inverse correlation between HuR expression and viral load and implicates HuR in cytokine dysregulation. Notably, HuR promoted cap-independent translation of viral RNA during later stages of infection, when cap-dependent translation is suppressed. These findings reveal a dual role for HuR: restricting viral RNA replication while enhancing translation, highlighting its critical, phase-specific function in the DENV life cycle.

**Author summary:** Understanding how Dengue virus interacts with human cells is key to finding better treatments. Our study looked at a human protein called HuR, which normally control the stability and the use of RNA inside our cells. We discovered that HuR has two important but opposite roles during dengue infection. Early on, it slows down the virus’s ability to make copies of its genetic material, helping to reduce infection. But later, when the virus struggles to use the normal method of making proteins, HuR steps in to help the virus make its proteins in a different way. This double-edged role of HuR shows how the virus cleverly uses host cell machinery at different stages of infection. Overall, this study uncovers the novel role of a specific RNA-binding protein of the host in orchestrating dengue virus infection and pathogenesis, highlighting its potential as a target for antiviral therapies.

**Graphical Abstract:** 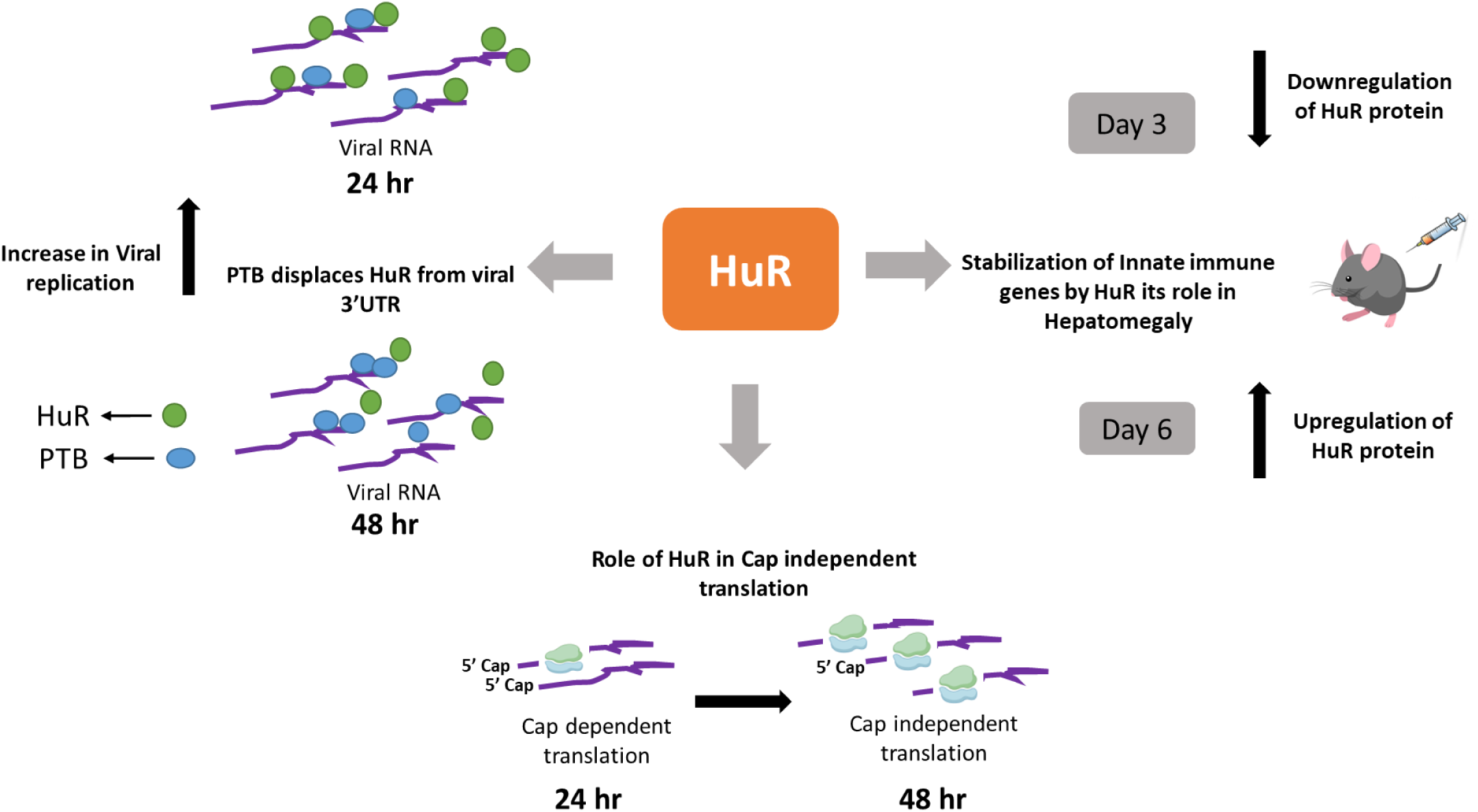

## Introduction

Dengue virus (DENV) is a mosquito-borne flavivirus belonging to the family *Flaviviridae*. DENV severe infection leads to dengue shock syndrome (DSS) in humans[1], causing 20,000 to 40,000 deaths per year globally, as per the World Health Organization [2]. Four antigenically different dengue virus (DENV) serotypes exist (DENV-1, 2, 3, and 4) with varying pathogenesis [3, 4]. The virus enters the host cells via receptor-mediated endocytosis and interacts with the endoplasmic reticulum (ER) membrane, forming replication complexes known as detergent-resistant membranes (DRMs) [5, 6]. These replication complexes have been shown to comprise both host and viral proteins, which interact with the viral RNA and affect its life cycle. Viral proteins such as NS4A and NS4B have been shown to remodel the ER membrane complex and aid in forming a virus replication complex. Although total RNA-binding proteins (RBPs) interacting with the viral untranslated regions (UTRs) have been reported earlier, no studies have been done to identify all the host proteins present in the viral replication complex during infection.

Although the viral RNA is capped, the viral 5’UTR has also been shown to translate via a cap-independent mechanism[7]. However, it is not yet understood when and how this cap-independent mechanism operates in the context of DENV infection, or which host factors are involved. RBPs such as La, polypyrimidine tract-binding protein (PTB), and HnRNP C1/C2 have previously been shown to interact with the viral RNA and play a role in virus translation and/or replication [8–10]. RBP binding may impact the interaction of viral UTR with other viral or host cellular factors, including proteins and miRNAs [11, 12]. It may also impact the circularization of the viral RNA, which is essential for virus replication. RBPs, such as Human antigen R (HuR) and La, have often been found to be associated with the viral replication complex as they aid in the regulation of host immune response genes [13, 14]. Utilization of these proteins by the virus during its life cycle also serves as a mechanism for suppressing the host’s immune response upon virus infection.

HuR is a vital RBP that has previously been shown to shuttle from the nucleus to the cytoplasm in the context of virus infection and has been shown to interact and stabilize many innate immune-responsive genes such as *IL-8, IL-6*, and *TNF alpha* [15], which have been shown to affect the cytokine pathway, which is central to dengue virus pathogenesis[16, 17] . HuR is also differentially regulated by many viruses, such as HCV, Zika virus, and CHV, and its modulation is known to directly affect virus-induced pathogenesis. However, the role of HuR protein in the DENV life cycle and pathogenesis has not been investigated. Given that HuR affects the replication of various viruses (HCV, Zika, and CVB3) through interaction with UTRs, we examined the interaction of HuR with DENV UTR [11, 18, 19]. Here, we report that HuR interacts with the 3’UTR of DENV and negatively regulates the viral life cycle by affecting the binding of PTB, which is known to positively influence viral replication [9]. Moreover, HuR was found to play an important role in cap-independent translation during the later stages of infection. Additionally, DENV–mediated regulation of HuR protein levels directly influences virus-induced pathogenesis by stabilizing host immunomodulatory genes. Overall, our work highlights the role of an important host factor, HuR, in viral replication, cap-independent translation, and pathogenesis.

## Results

1. **HuR protein is associated with the viral replication complex –** Understanding the composition of the Detergent Resistant Membranes (DRMs), which comprise the viral replication complex, is crucial in dissecting the regulation of viral replication inside the host. Therefore, DRMs were isolated from virus-infected Huh7 cells at 48 h post-infection by treating the cell lysate with and without detergent, followed by ultracentrifugation (Fig. 1A). The results showed a significant shift in the position of some known RBPs, such as HuR and PTB, toward a lower fraction along with the viral NS5 protein, which served as a positive control, suggesting their presence in DRM fractions. Semiquantitative PCR showed the presence of viral negative strands in the 2nd, 3rd, and 4th fractions, which also confirmed the viral replication complex (Fig. 1B). Mass spectrometry data of fractions 2, 3, and 4 identified a lot of RBPs such as HuR and PTB, along with heat shock proteins and chaperons, all of which were found to be associated with the viral replication complex and could play critical role in the viral life cycle (Supplementary Table 1). The viral NS5 protein is predominantly nuclear, and the cytoplasmic localization is within the viral replication complex (Supplementary Fig. 1 (A and B)). So NS5B was used as a marker to determine the site of virus replication. Its cytoplasmic localization was visualized using a cytoplasmic-specific permeabilization agent, digitonin. To check whether the RNA binding protein HuR co-localizes with the viral NS5 protein in the DRM’s, confocal microscopy was performed at 48 h post-infection using digitonin-mediated permeabilization. PTB was used as a positive control, which was previously shown to aid in the viral life cycle (Fig. 1E). The data analyses showed that HuR and PTB significantly colocalized with the viral NS5 protein with a colocalization coefficient of >0.5 (Fig. 1F). Results showed, both HuR and PTB are present in the viral replication complex and may interact with the viral RNA.
2. **HuR negatively regulates Dengue replication** – To assess the effect of virus infection on the overall levels of HuR and PTB, infection was carried out at (MOI of 1 and 2) in Huh7 cells, followed by reverse transcription polymerase chain reaction (RT-PCR) to check for viral RNA levels (Fig. 2A and B). A steep increase in the positive and negative strands of the viral RNA was observed from 24 to 48 h post-infection, which represents a high rate of viral RNA replication during this period. To investigate whether the levels of PTB and HuR changes in this active phase of viral RNA replication, western blotting was done at 24 h and 48 h post-infection (Fig. 2C). Interestingly, a significant increase in the levels of HuR and PTB was observed at 24 h, followed by a sharp decrease at 48 h, which coincides with the decrease in host translational machinery as observed in dengue infection[20]. However, the increase in HuR levels was much more pronounced than that of PTB at 24 h post-infection (Fig. 2C). Next, to determine further whether this change in protein level regulation is due to the shuttling of these proteins from the nucleus to the cytoplasm post-infection, we measured the intensity of these proteins at a single infected cell resolution (Supplementary Fig 2). No significant difference was seen in the abundance of HuR in the cytoplasm upon virus infection, indicating no relocalization of the protein to the cytoplasm post-infection. To evaluate the role of HuR in the viral life cycle, HuR was silenced using siRNA targeting HuR mRNA in Huh7 cells, followed by virus infection (Fig 2D and E). Upon partial silencing of HuR, a significant upregulation of the viral RNA was observed at 48 h and 72h post-infection. In contrast, overexpression of HuR led to a significant downregulation of viral RNA levels, post-infection (Fig. 2F and G). Furthermore, to determine whether HuR influences the viral RNA translation or replication, we used a dengue sub-genomic replicon construct containing a luciferase reporter gene in place of the structural proteins (Fig. 2H). The luciferase values did not show significant changes upon silencing of the *HuR* gene (Fig. 2I and J), suggesting that HuR predominantly affects the viral RNA replication via interacting with the viral 3’UTR, as shown in Fig. 3. Further, to evaluate the impact of total knockout (KO) of HuR and PTB on the viral life cycle, CRISPR-mediated KO of both HuR and PTB was performed in HeK293T cells (Supplementary Fig. 3). As anticipated, viral replication was significantly upregulated (by approximately 100%) in HuR KO cells, whereas the PTB KO resulted significant downregulation (by approximately 80%) of viral replication (Supplementary Fig. 3). Overall, these results suggest that HuR acts as a negative regulator of viral RNA replication, in contrast to PTB, which has been reported as a positive regulator of viral RNA replication[9].
3. **HuR protein interacts with the viral 3’UTR –** To determine whether HuR protein interacts with viral RNA, a direct UV-crosslinking experiment was performed using recombinant HuR protein and radiolabelled viral 5’ and 3’ UTR RNA, followed by RNase treatment and autoradiography. The results showed a significant interaction of HuR protein with the viral 3′UTR that increased with increasing protein concentration (Fig. 3A). To further confirm the specificity of this interaction, a competitive UV-crosslinking experiment was conducted in the presence of self and non-self-cold RNAs. The results indicated a specific interaction of HuR protein with the viral 3’UTR, but not with the viral 5’UTR (Fig. 3B and C). Additionally, to validate the specificity of the interaction, competitive UV-crosslinking experiment was carried out using S10 lysates of Huh7-infected cells. A specific decrease in the intensity of a band corresponding to the molecular weight of HuR was observed, which was further confirmed via immunoprecipitation with anti-HuR antibody post UV-crosslinking (Fig. 3D and E). Further, to examine the interaction of HuR protein with the viral RNA following DENV 2 infection, RT-PCR was performed post-immunoprecipitation with anti-HuR antibody. Results showed that HuR protein does interact with the viral RNA post-infection (Fig. 3F).
4. **PTB displaces HuR from the 3’UTR of the viral RNA, leading to an increase in viral RNA levels upon infection –** To understand the mechanism by which HuR regulates the viral life cycle, the binding site of HuR protein at the 3’UTR of the viral RNA was investigated using CatRapid tool and sequence-specific interaction (Supplementary Fig. 4A). Interestingly, the binding site for PTB was found to overlap with the binding site for HuR protein (Supplementary Fig. 4A). To assess the interplay between these two proteins, immunoprecipitation was performed using anti-HuR and anti-PTB antibodies, and the associated viral RNA was quantified by RT-PCR at different time intervals post-infection. HuR was more strongly associated with viral RNA at 24 h, while PTB association was greater at 48 h (Fig. 4A - D). These results correlate with our earlier results showing higher upregulation of HuR compared to PTB at 24 h post-infection (Fig. 2). Additionally, PTB was found to displace HuR protein from the 3’UTR of the viral RNA, whereas HuR protein could not displace PTB in the in vitro UV-crosslinking assay (Fig. 4E). Overexpression of PTB displaced HuR from the viral RNA in IP-RT (immunoprecipitation followed by reverse transcriptase polymerase chain reaction), whereas HuR overexpression failed to displace PTB (Fig. 4F-I). To further assess HuR displacement by PTB, the putative binding sites of HuR and PTB were mutated in the 5′Luc3′ reporter construct (Fig. 4J). The wild-type and mutant 5’Luc3’ reporter DNAs were transfected into Huh7 cells, and luciferase activity was measured at 48 h. The HuR binding mutant showed higher luciferase activity compared to the PTB binding mutant (Fig. 4K). To confirm the impact of these mutations on protein binding, IP-RT was performed after transfection with wild-type and mutant constructs (Supplementary Figure. 4B-E). The HuR binding mutant showed greater PTB interaction, while the PTB binding mutant exhibited greater HuR interaction with the reporter construct, reconfirming the interplay between HuR and PTB at the viral 3′UTR.
5. **HuR positively regulates cap-independent translation at 48 h post-infection –** To understand the role of HuR in cap-independent translation of dengue virus, reporter constructs 5’UTR Luc and 5’UTR-Luc-3’UTR were transfected into Huh7 cells in the background of virus infection at MOI-1. Rluc was used as an internal control for cap-dependent translation. An increase in Fluc/Rluc ratio was observed at 24 h and 48 h post-infection in cells transfected with 5’UTR-Luc-3’UTR but not in 5’UTR-Luc (Fig. 5A-C). Results clearly show cap-independent translation at a later time point post-infection, and the specific role of 3’UTR in its regulation[7], along with a significant decrease in cap-dependent translation, which is further evidenced by Rluc activity and polysome analysis at 48hr post-infection (Supplementary Fig. 5G). Further, to specifically assess the role of HuR in cap-independent translation at later time points, 5’UTR Luc and 5’UTR-Luc-3’UTR reporter constructs were transfected into HEK293T HuR KO cells, and HEK293T was used for comparison. The Fluc/Rluc ratio in HuR KO cells was found to be significantly downregulated at 48hr as compared to normal cells (Fig. 5C-D). The absolute values are shown in (supplementary table 2). To reconfirm the role of HuR in cap-independent translation of dengue viral RNA, the reporter constructs were transfected in the presence of LY294002, a known inhibitor of cap-dependent translation. HuR KO cells showed a significant decrease in cap-independent translation activity of DENV at 48 hr post-transfection (Fig. 5F-G). As expected, the HuR binding mutant showed a significant decrease in Fluc/Rluc ratio in the background of virus infection, even at 48 hr hours post-infection when the HuR levels were found to be significantly downregulated (Supplementary Fig. 5H) and (Fig. 2C).
6. **HuR protein levels inversely correlate with viral RNA levels and its mRNA targets during infection. -** To assess HuR protein levels upon infection and its role in virus-induced pathogenesis, we checked the levels of HuR protein in AG129 mice in the brain and liver tissues at day 3 and day 6 post-DENV infection. HuR protein levels were downregulated at day 3 post-infection and upregulated at day 6 post-infection in mouse liver tissues (Fig. 6B-E). The levels of HuR protein were further found to inversely correlate with the viral RNA levels in liver tissues at Day 3 and Day 6 post-infection (Fig. 6F), showing a negative regulation of dengue virus by HuR protein. The mRNA levels of the important immunoregulatory pathways, such as IL-8, IL-6, and TNF-alpha, which are known to be stabilized or destabilized by HuR protein[21], were checked (Fig. 6G), the levels of IL-8 and TNF-alpha were found to be upregulated in both mouse liver tissue samples and HuR KO cells upon dengue infection (Fig. 6H-I) and (Supplementary Fig. 6). The association of TNF-alpha and IL-8 mRNA with HuR protein was further found to be downregulated upon virus infection at day 3 post-infection (Fig. 6J-M), thereby highlighting the role of HuR protein as an important player in pathogenesis via the negative regulation of TNF-alpha and IL-8 mRNAs upon dengue infection.

## Discussion

DENV replication in the human host begins with the Langerhans cell beneath the skin, followed by infection in the liver, the predominant site of virus replication and pathogenesis [22–24]. Virus replication inside the host cells is affected by various host factors, including RBPs and miRNAs that bind to the viral UTRs and may act as proviral or antiviral factors. An abundance of these factors ultimately decides the virus’s tissue specificity in the human host cells [12, 25] or mosquito cells [26]. Upon entry into the cells, the virus forms replication complexes with the ER membrane, which is known to contain many host and viral proteins. The replication complexes of other flaviviruses such as HCV and Zika, have been characterized previously, showing many RBPs such as La and PTB, and chaperons such as HSPs (Heat Shock proteins). Our mass spectrometry data of the DENV replication complexes also showed a similar profile with many hnRNPs, HSPs, and RBPs present in the virus replication complex [27–29]. We also observed the enrichment of an important RBP, HuR, in our DRM fraction using western blotting. However, our mass spectrometry data did not detect the specific presence of HuR protein in the spectra, which may be due to the presence of HuR protein in small amounts in the entire cytoplasmic fractions, as evidenced by western blotting.

**Figure 1.**
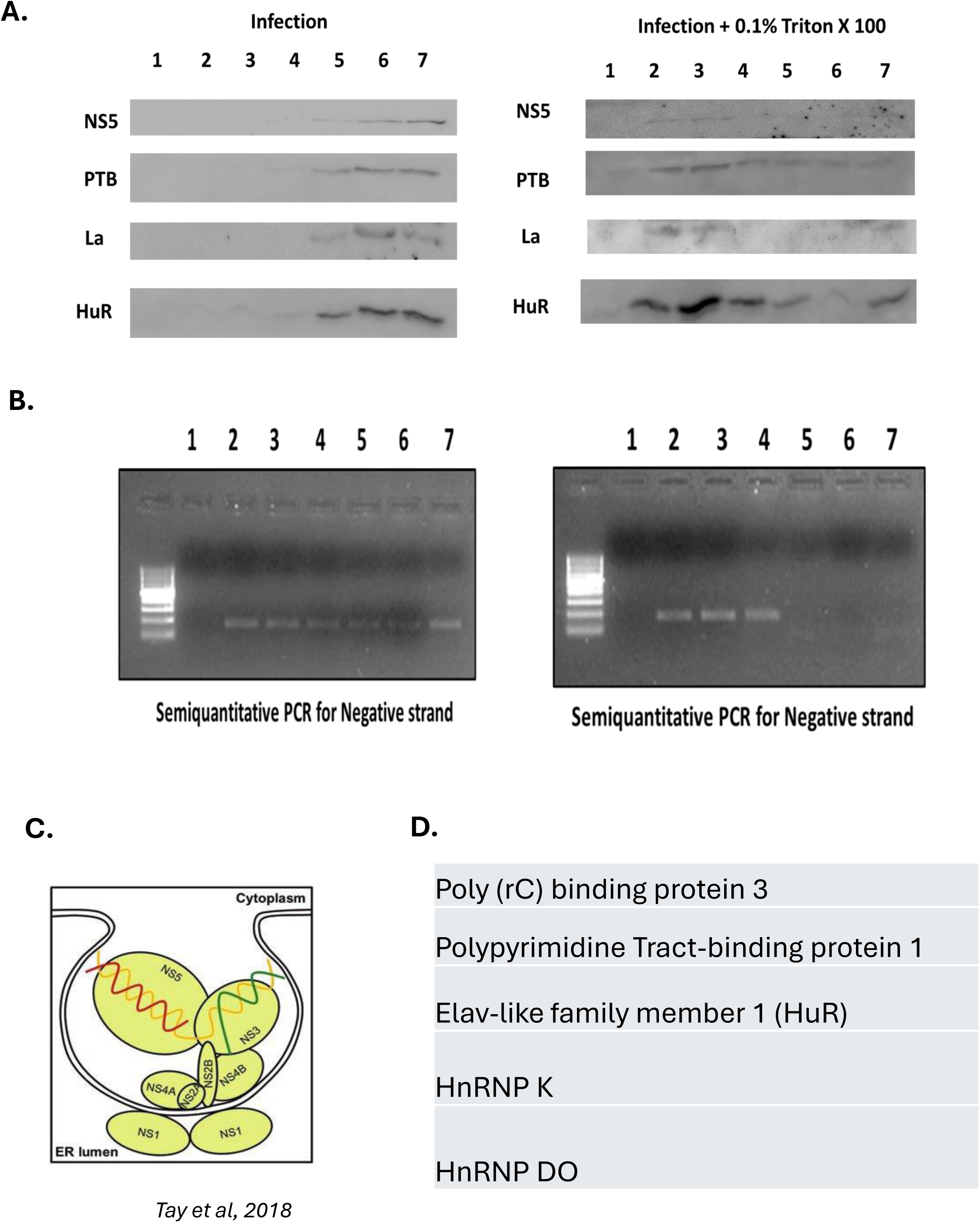
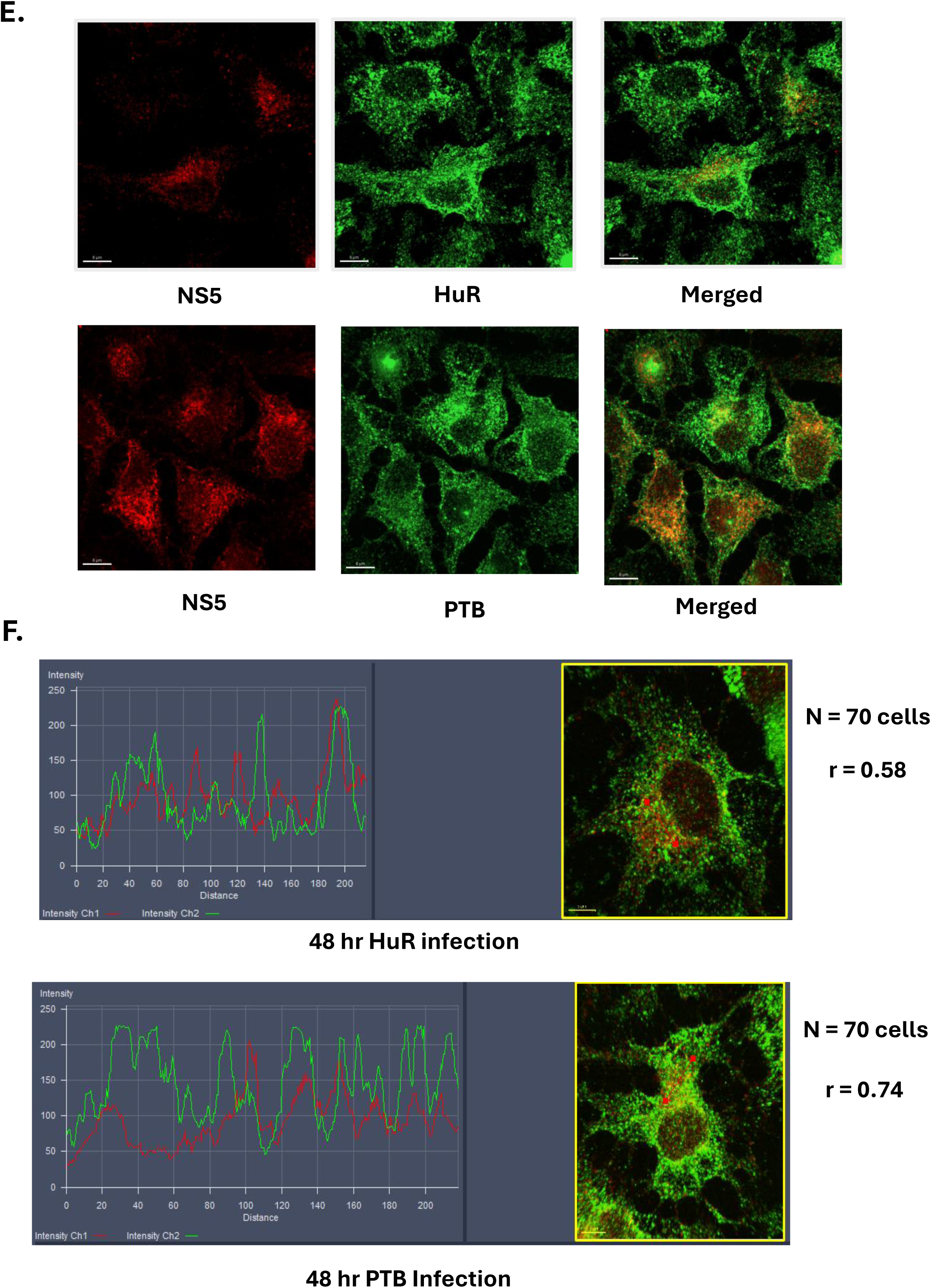
HuR protein is associated with the viral replication complex. (A) Isolation of DRMs using sucrose density centrifugation and Western blot to check for the presence of Host and Viral proteins. (B) Semiquantitative PCR shows enrichment of negative strands in fractions 2,3, and 4. The negative strand of the viral RNA was checked with the primers against NS5 region of the viral genome. (C) Localization of NS5 in the cytoplasm, adapted from Tay et al, 2018. (D) Mass spectrometry data showing important host proteins. The fractions 2,3, and 4 were isolated and sent for Mass spectrometry. A complete list of host proteins identified has been separately provided in Supplementary Table 1 (E, F) Co-localization of HuR and PTB proteins in the cytoplasm with a co-localization coefficient. The virus was infected with MOI=1 in Huh 7 cells, followed by detergent-mediated permeabilization with digitonin. HuR, NS5, and PTB were stained with appropriate secondary antibodies for visualization. The co-localization analysis is from 60 cells and the co-localization analysis was done using Image J software.

**Figure 2.**
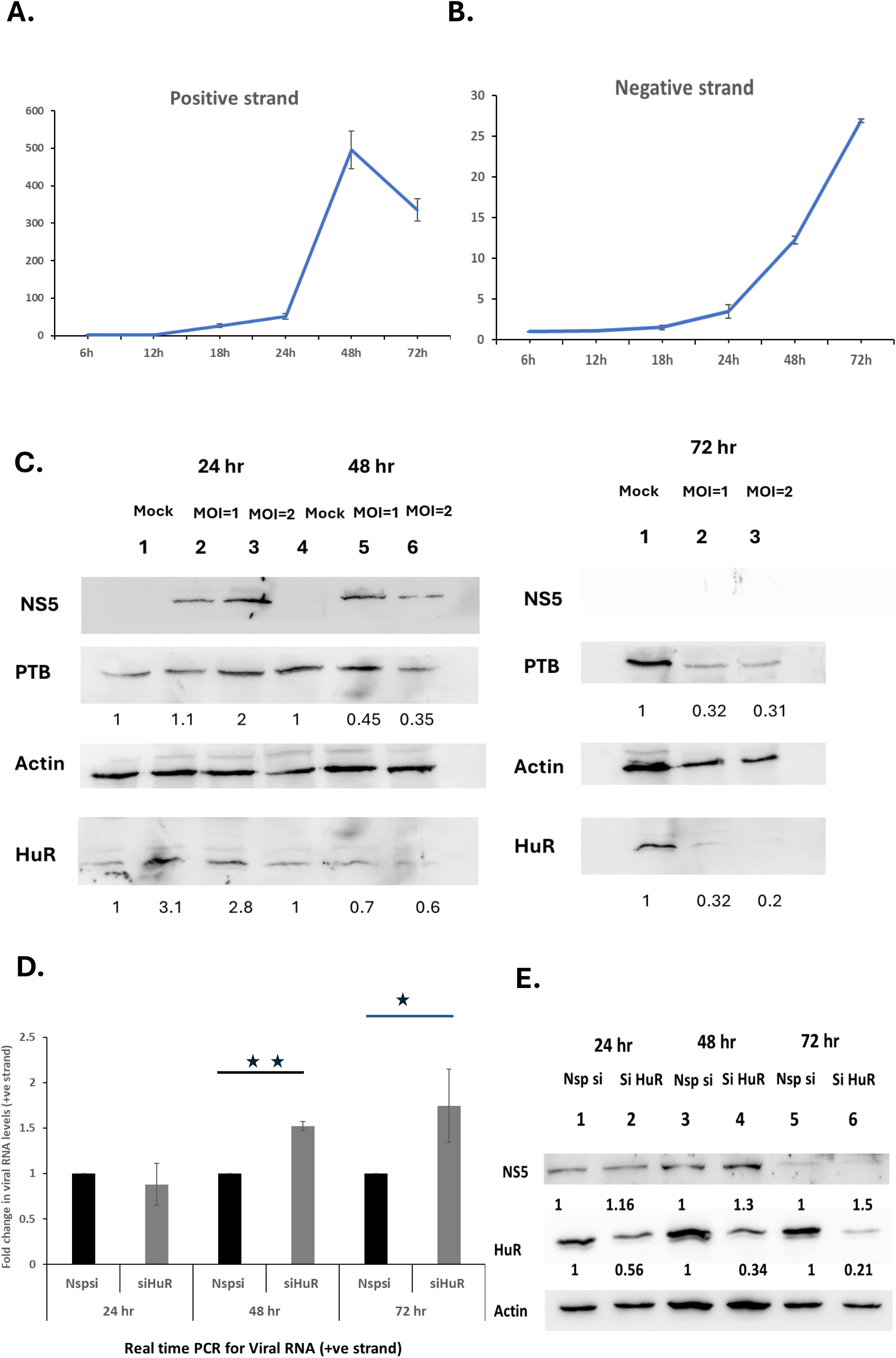
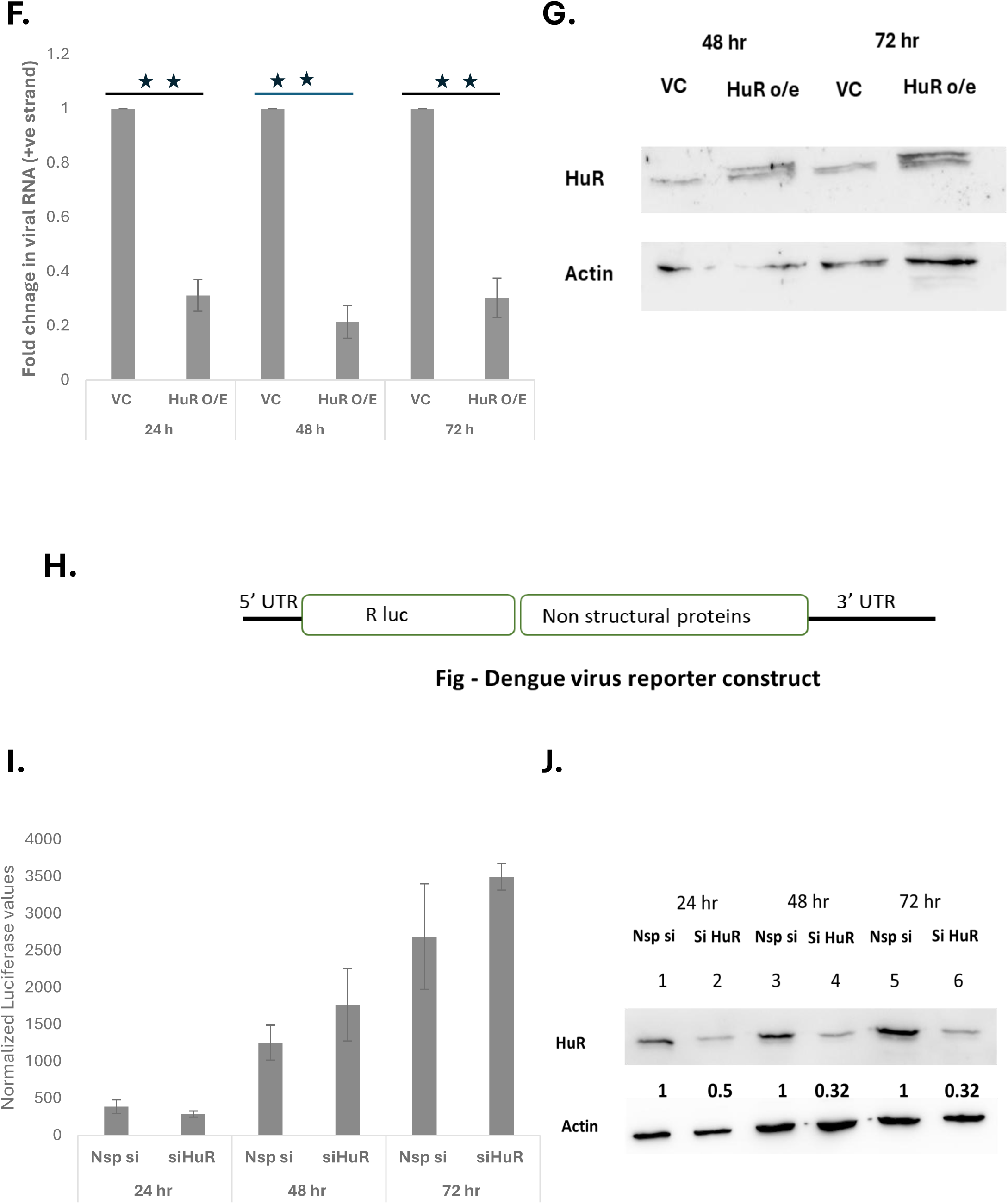
HuR negatively regulates Dengue replication. (A, B) - Dengue growth curve at MOI = 1. Virus infection was given at MOI = 1 in Huh 7 cells, followed by RNA isolation and Real Time PCR against the viral genome at different time points. (C) - HuR and PTB protein levels during infection in Huh 7 cells. Post virus infection proteins were isolated, and Western blotting was done to check the levels of the proteins with Actin as an internal control. (D, E) - siHuR leads to an increase in viral RNA levels post-infection. HuR protein was silenced with 150nM of siRNA against HuR mRNA, followed by infection after 16 hours. (F, G) - HuR over-expression leads to a decrease in viral RNA levels post-infection. HuR over- expression was done with 100ng of PCDNA3-HuR, and PCDNA3 was taken as a vector control, followed by infection with DENV Serotype 2 at MOI=1. (H) - Schematics of sub-genomic replicon. (I, J) - siHUR has no effect on the luciferase value of the luciferase reporter construct upon transfection. 150nM of siRNA against HuR mRNA was transfected followed by transfection of 500ng of Dengue SgDV-R2A RNA. The Luc values were normalized with the total protein concentration. Students t-test was used for statistical analysis. P<0.05 = *, P<0.01 = **

**Figure 3.**
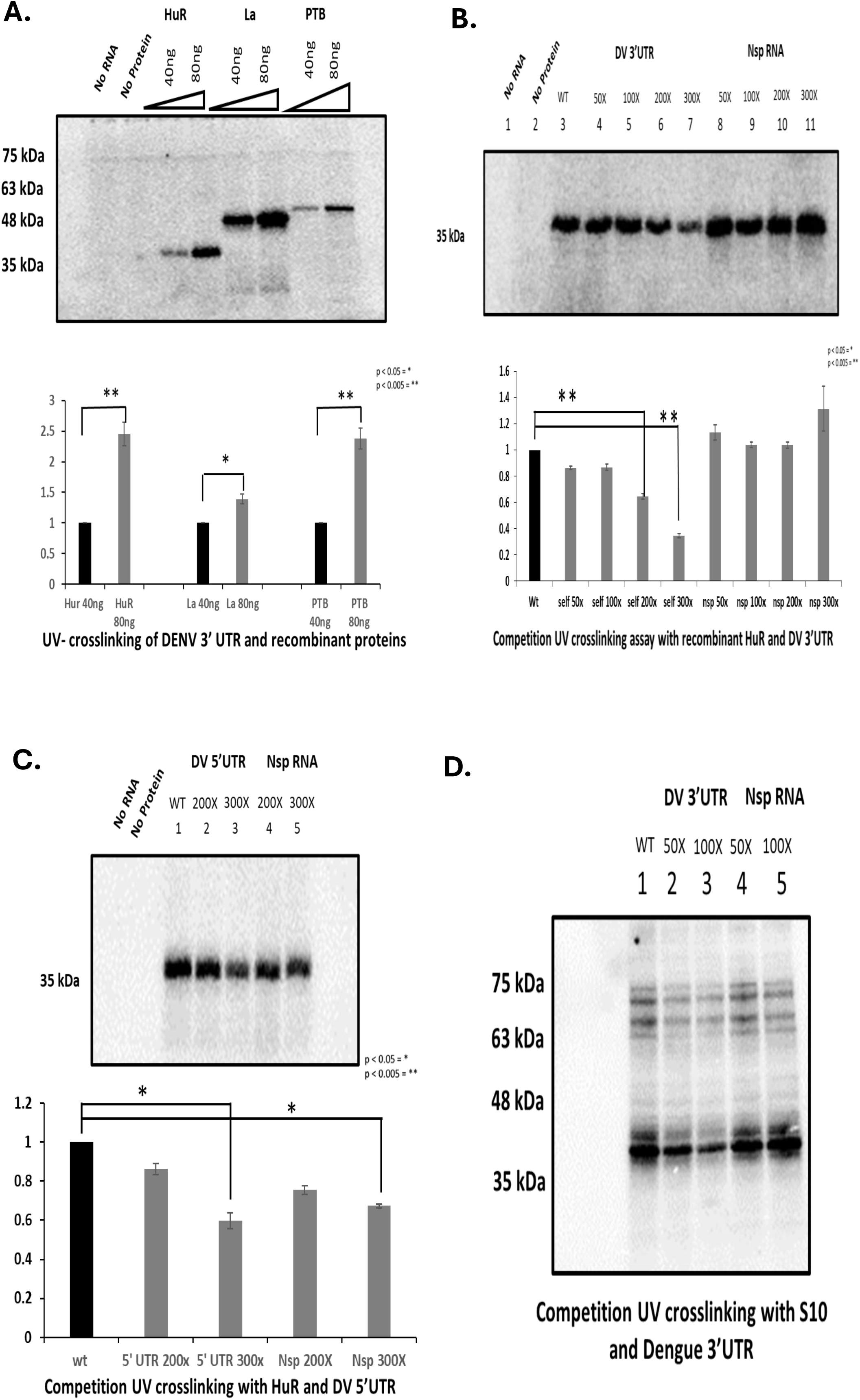
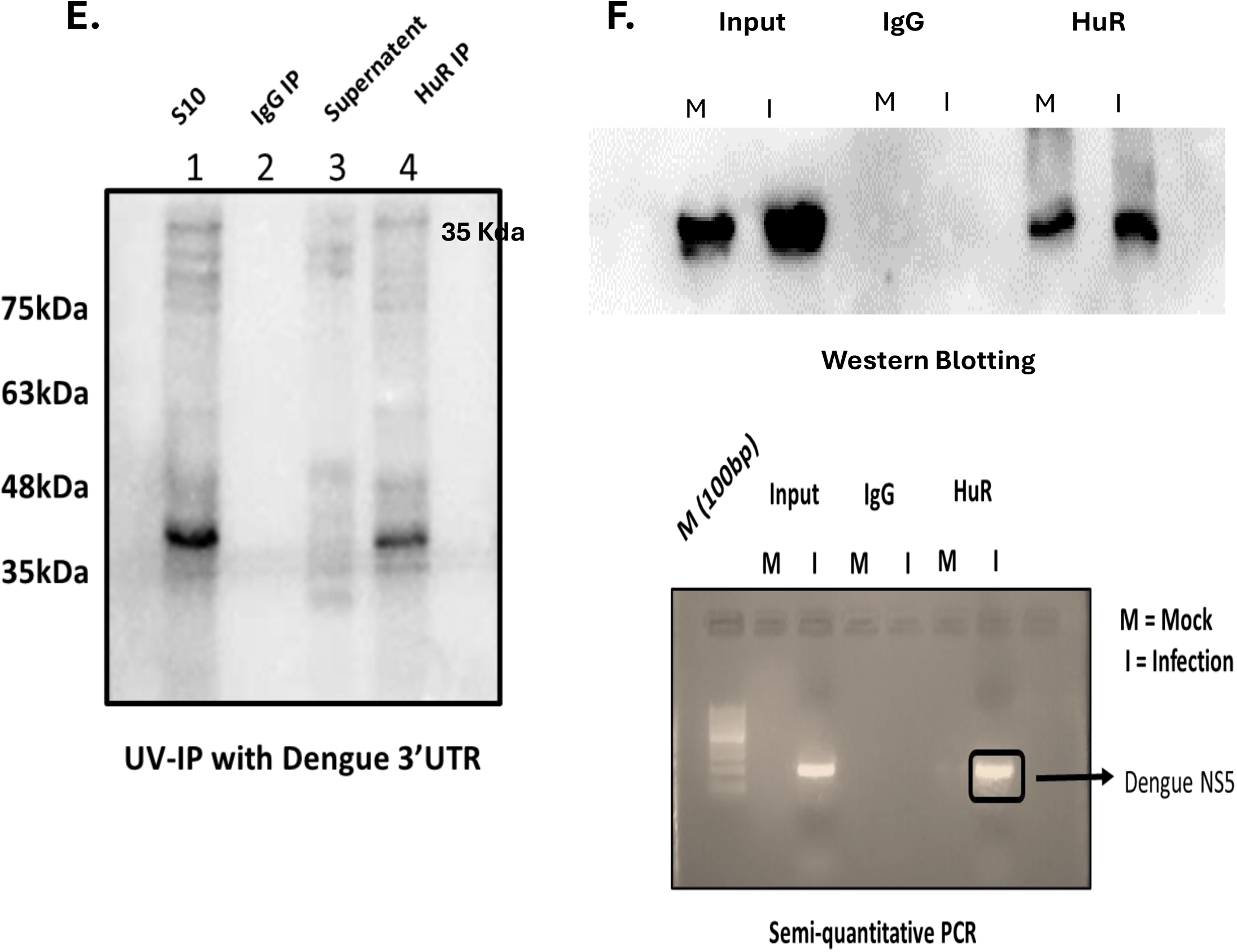
HuR protein interacts with Dengue virus 3’UTR. (A) - UV cross-linking assay showing the interaction of HuR and PTB with Dengue virus 3’UTR. (B) - Competition UV crosslinking assay showing high affinity of interaction of the viral 3’UTR. (C) - Competition UV crosslinking assay showing low affinity with viral 5’UTR. (D) - Competition UV crosslinking with S10 extract showing specific interaction with the viral 3’UTR. (E) - UV-IP to confirm the identity of the band as HuR protein. (F) - IP-RT to show the interaction of HuR with the viral RNA upon viral infection in HuH 7 cells. Immunoprecipitation was done using anti-HuR antibody, followed by Real Time PCR for viral RNA. Students t-test was used for statistical analysis. P<0.05 = *, P<0.01 = **

**Figure 4.**
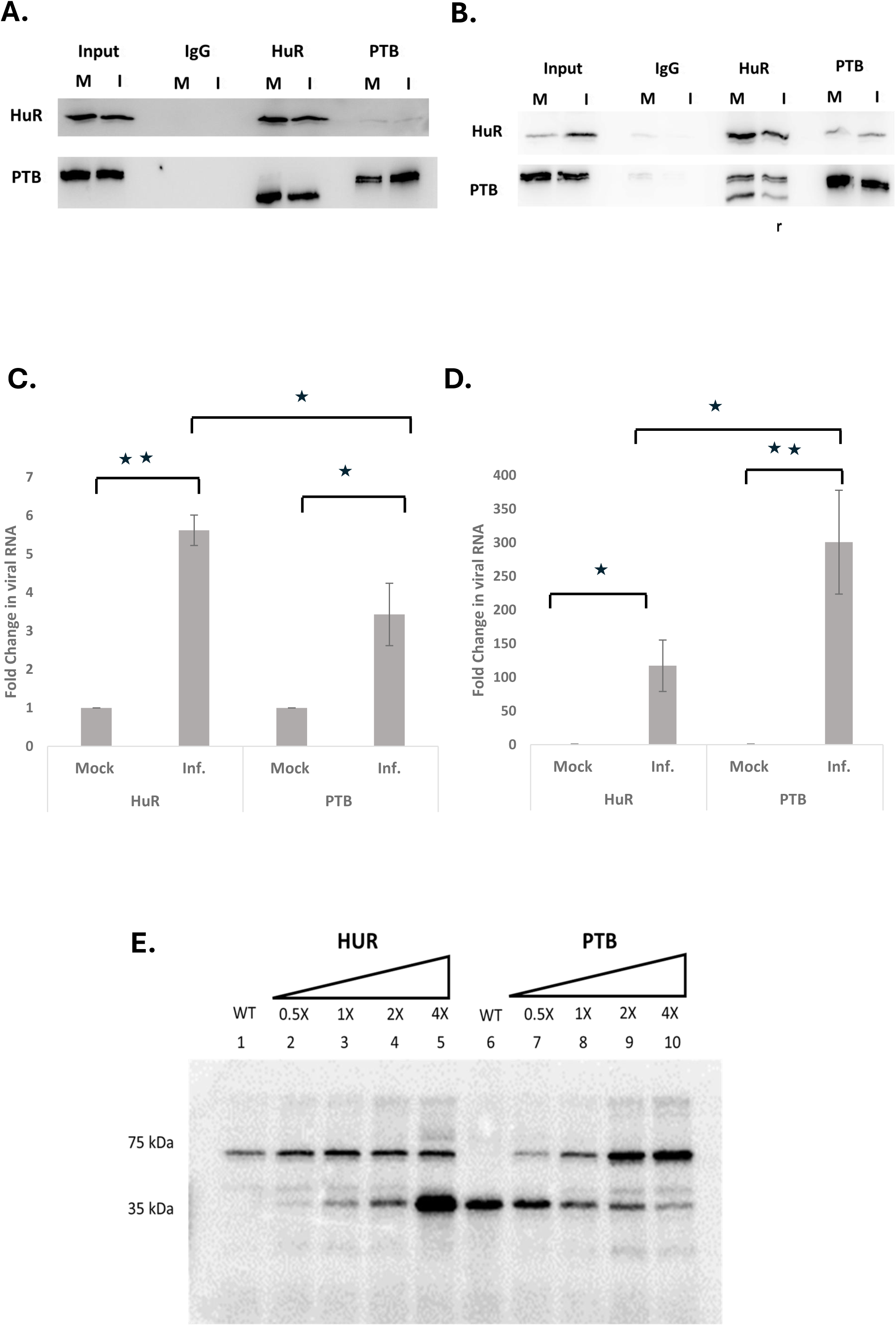
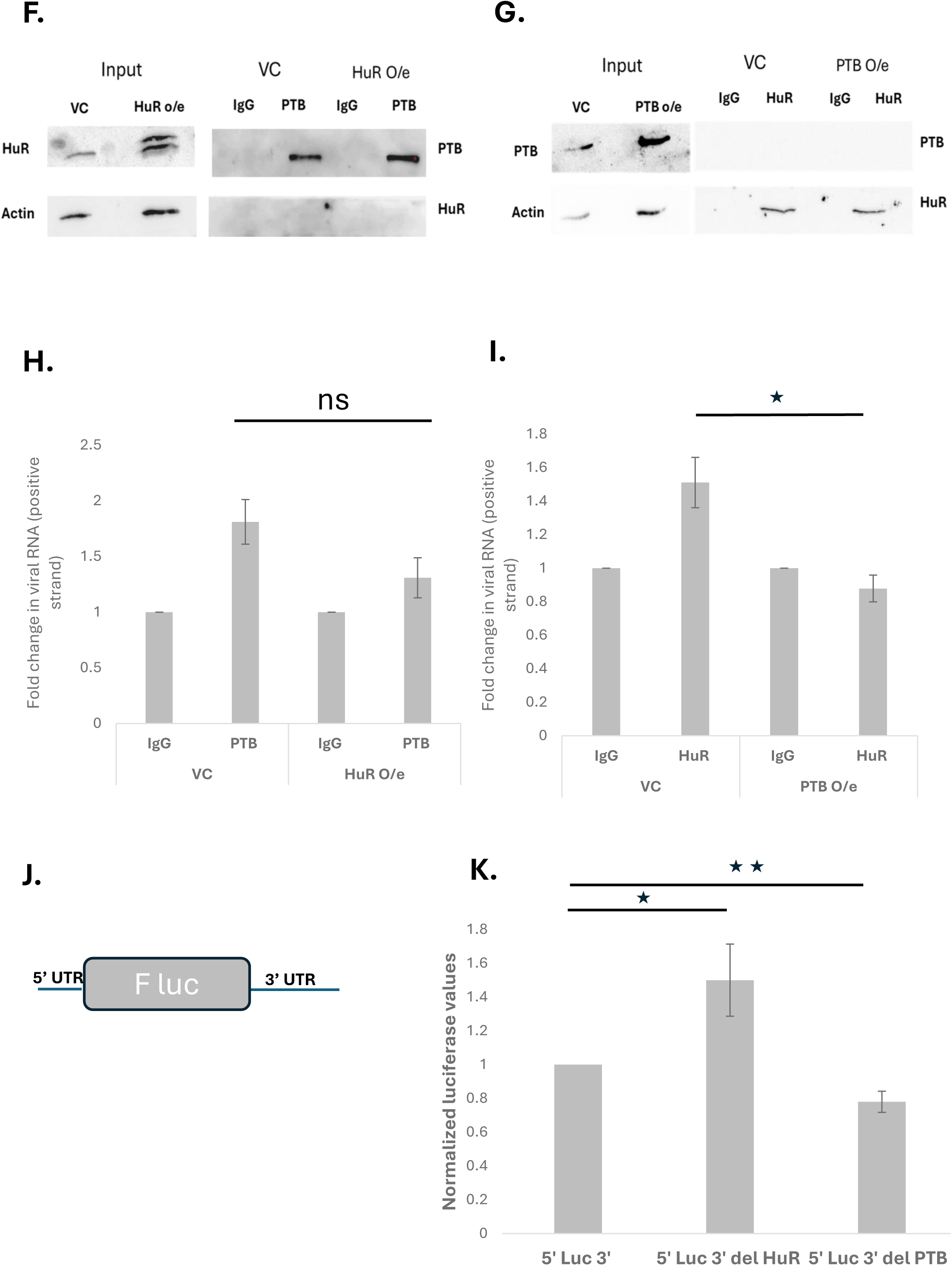
PTB displaces HuR from the 3’UTR of the viral RNA leading to an increase in viral RNA levels upon infection. (A, B) Western blotting to check for HuR and PTB upon immunoprecipitation with anti-HuR and anti-PTB antibodies at 24 hr and 48 hr, respectively. (C, D) – Real Time PCR of the RNA isolated from immunoprecipitated fraction showing the association of viral RNA at 24hr and 48 hr, respectively. Normalization was done with the Ct values of input RNA. (E) - Competition UV crosslinking showing PTB can displace HuR from viral 3’UTR. HuR and PTB recombinant proteins were added with increasing ratio to check for competition. (F, G) Western blotting to check for HuR and PTB upon immunoprecipitation with anti-HuR and anti-PTB antibodies at 24 hr and 48 hr, respectively, in the background of HuR and PTB over-expression. (H, I) – Real Time PCR of the RNA isolated from immunoprecipitated fraction showing the association of viral RNA at 24hr and 48 hr, respectively. Normalization was done with the Ct values of input RNA. (J, K) - Luciferase activity upon transfection of wild-type mutant reporter constructs. The predicted binding sites of HuR and PTB were mutated as shown in the supplementary figure 4. Students t-test was used for statistical analysis. P<0.05 = *, P<0.01 = **

**Figure 5.**
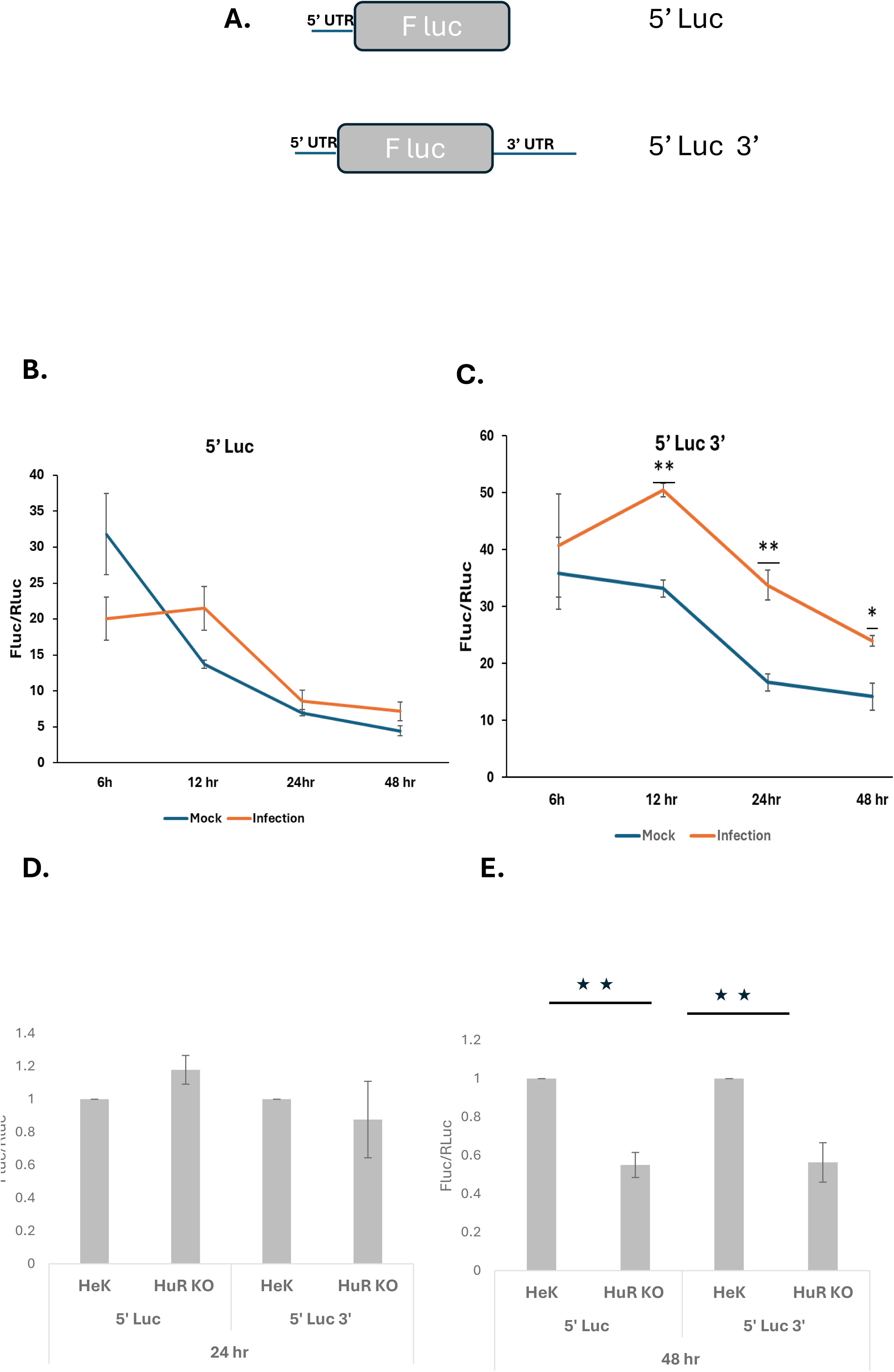
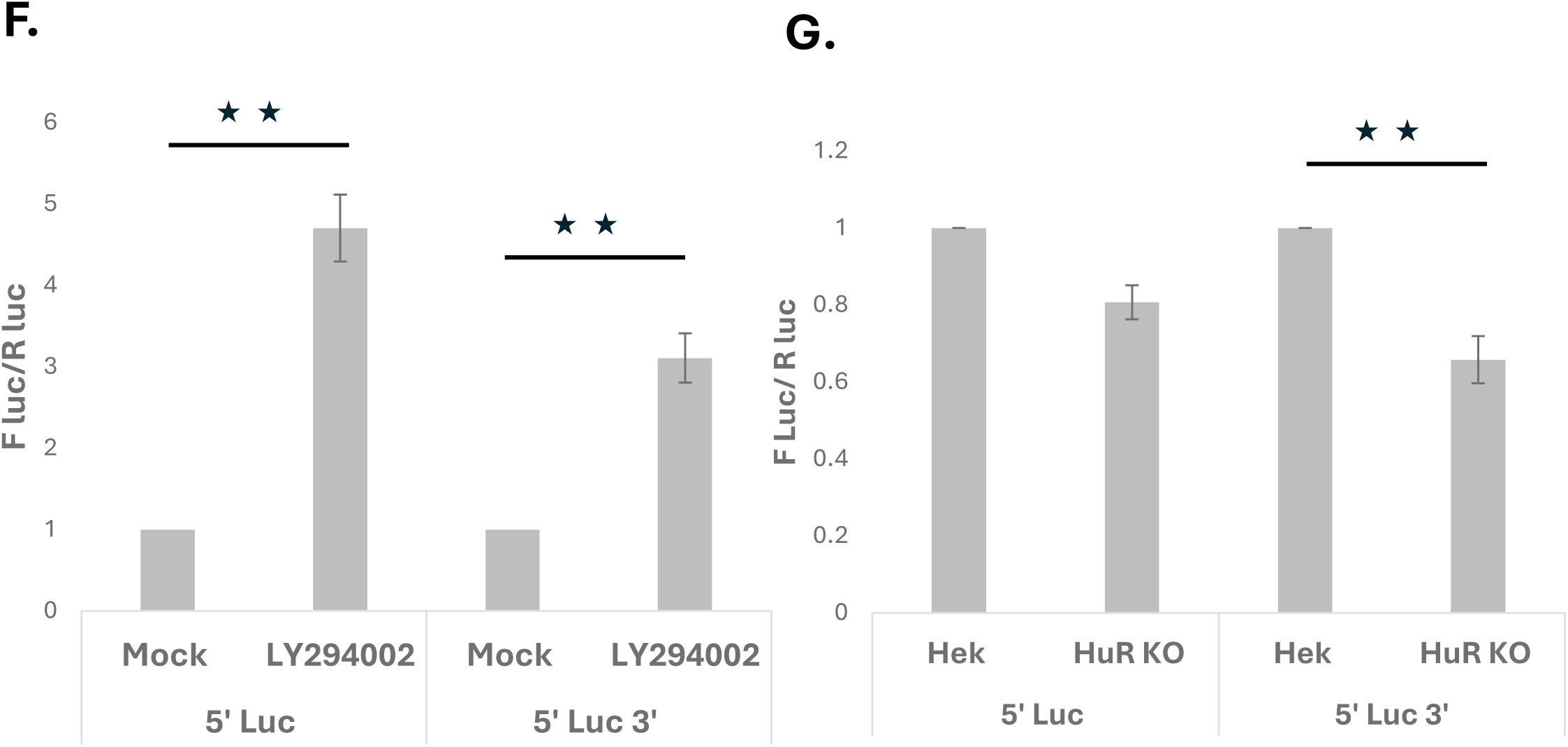
HuR positively regulates cap-independent translation at 48hr post infection. (A) - Schematic representation of dengue 5’Luc and 5’Luc 3’ constructs (B) - Fluc/Rluc activity of 5’ Luc at different time points in the background of infection at different time points in Huh 7 cells. Virus infection was given at MOI=1 post-transfection of 5’Luc reporter constructs, and PCDNA3-Rluc was taken as a transfection control. (C) - Fluc/Rluc activity of 5’ Luc 3’ at different time points in the background of infection at different time points in Huh 7 cells. Virus infection was given at MOI=1 post-transfection of 5’Luc 3’ reporter constructs, and PCDNA3-Rluc was taken as a transfection control. (D, E) - Fluc/Rluc activity at 24 hr and 48 hr post-transfection in HEK 293T and HEK 293T HuR KO cells. 5’Luc and 5’Luc3’ were transfected along with PCDNA3-Rluc as a transfection control, and the Luc values were taken post-harvesting at 24 hours and 48 hours, respectively. (F) - Fluc/Rluc at 48 hr post-transfection in the presence of cap inhibitor LY294002. 5’Luc and 5’Luc3’ were transfected along with PCDNA3-Rluc as a transfection control, and the Luc values were taken post-harvesting at 48 hours. (G) - Fluc/Rluc at 48 hr post-transfection of 5’Luc and 5’Luc3’ in the presence of cap inhibitor LY294002 in HEK 293T and HEK 293T HuR KO cells. 5’Luc and 5’Luc3’ were transfected along with PCDNA3-Rluc as a transfection control, and the Luc values were taken post- harvesting at 48 hours. Students t-test was used for statistical analysis. P<0.05 = *, P<0.01 = **

**Figure 6.**
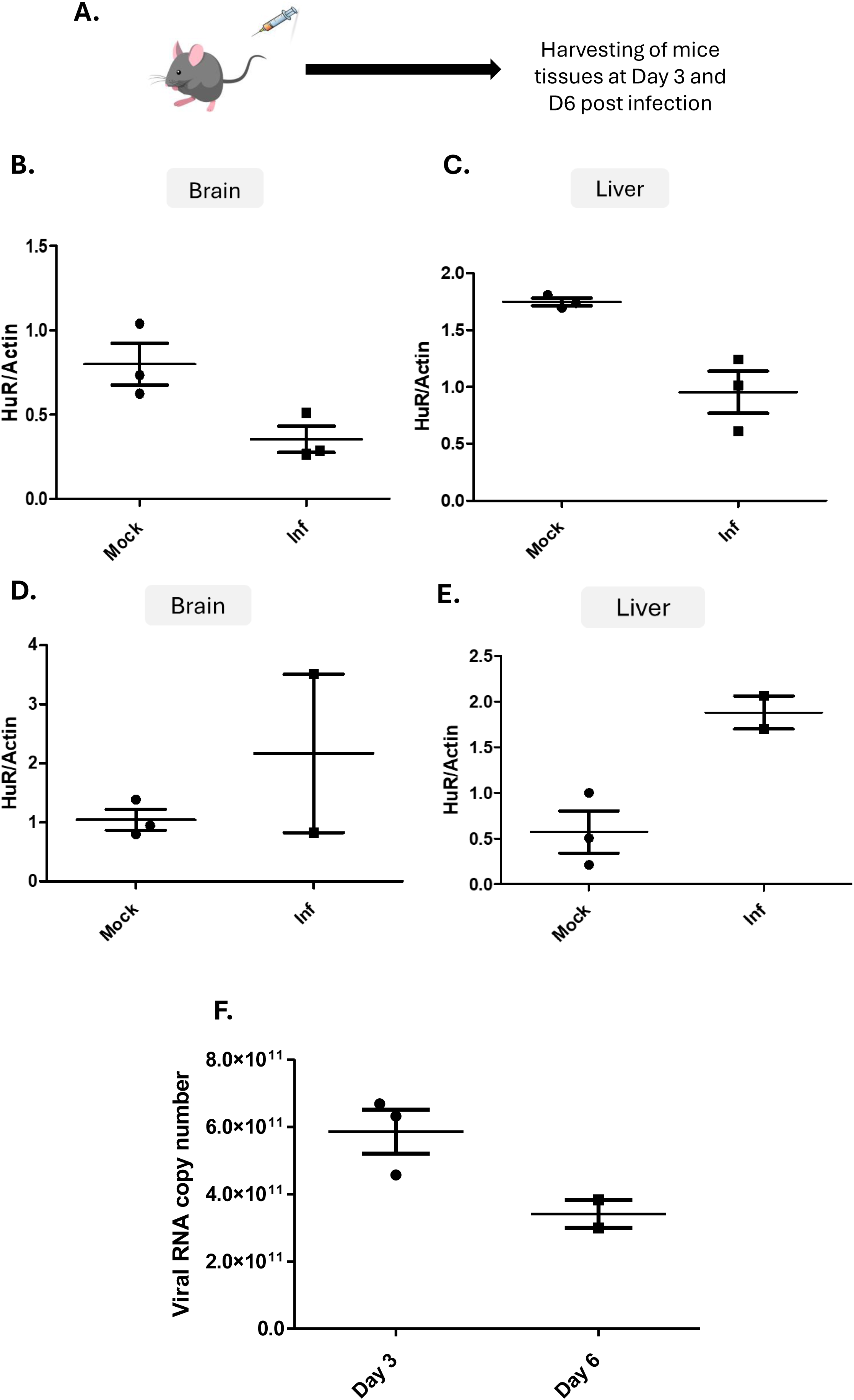
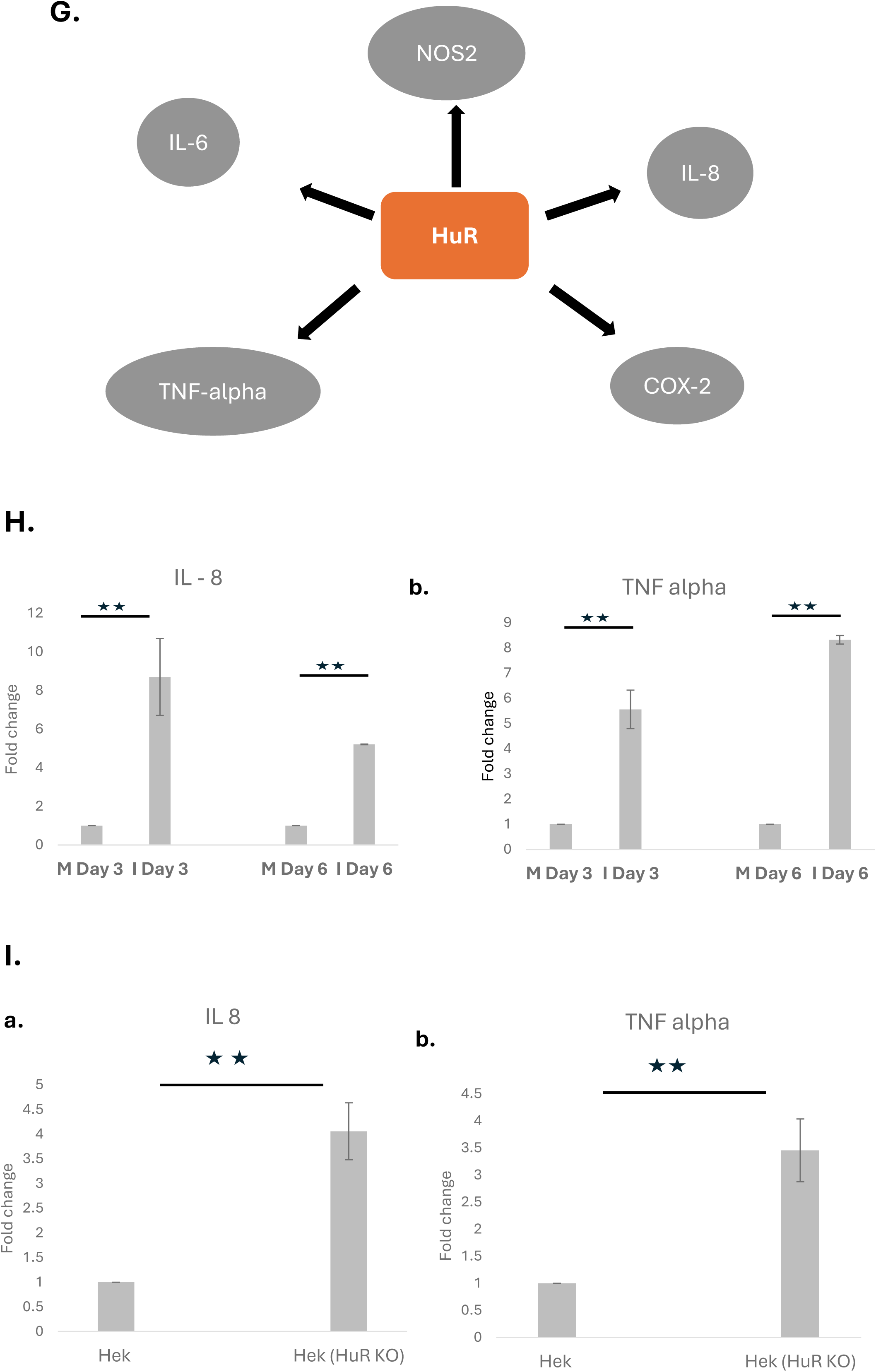
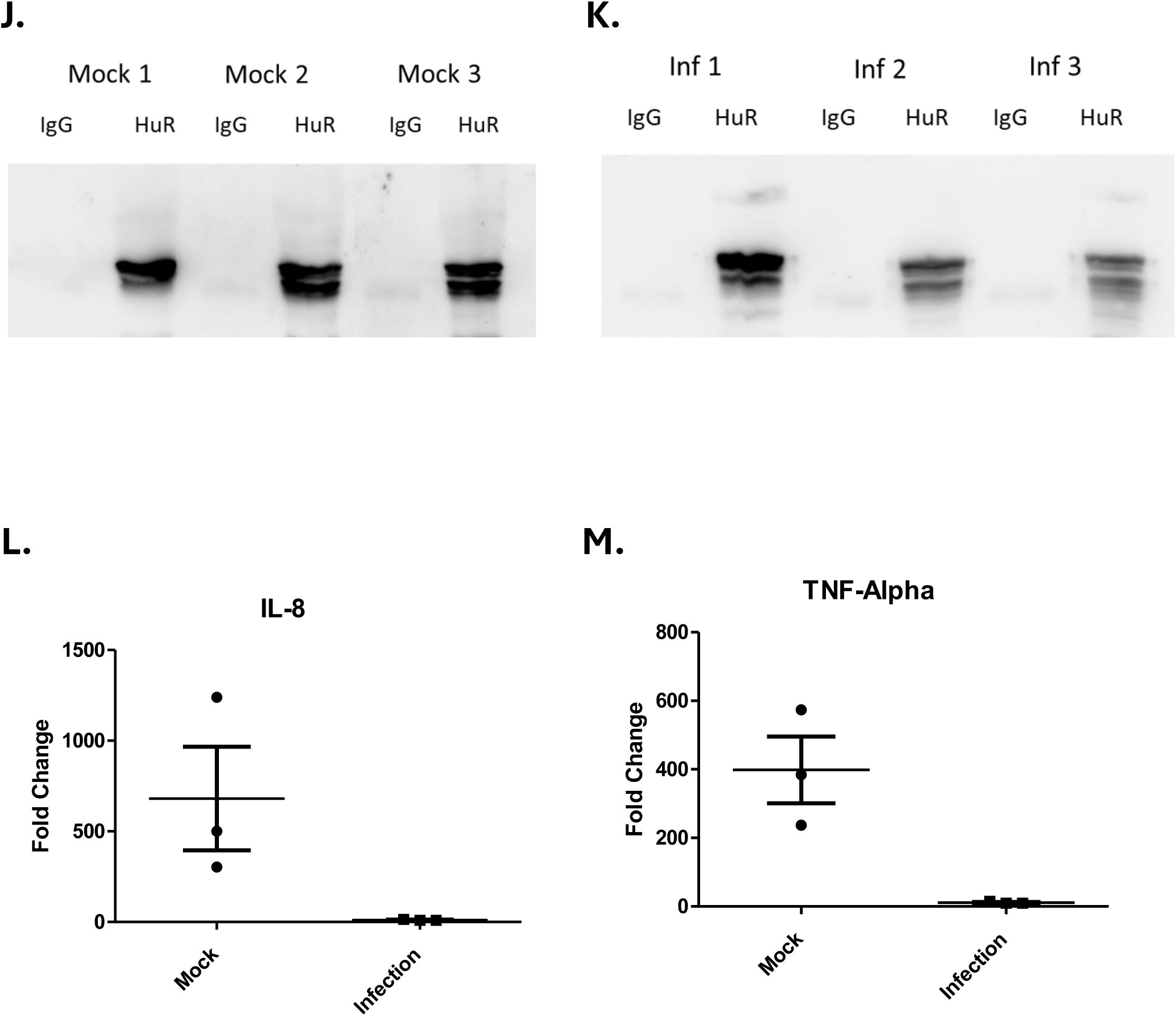
HuR protein levels inversely correlate with viral RNA levels and its mRNA targets during infection. (A) - Schematic showing virus infection in mice. (B, C) - HuR levels in the mouse brain and liver tissues at Day 3 post-infection. Virus infection was given in AG129 mice, followed by western blotting to check HuR levels. Actin was taken as an internal control for this experiment. (D, E) - HuR levels in the mouse brain and liver tissue at Day 6 post-infection. Virus infection was given in AG129 mice, followed by western blotting to check HuR levels. Actin was taken as an internal control for this experiment. (F) - Viral RNA copy number in liver tissue at Days 3 and 6 post-infections. Total RNA was isolated from infected mice liver tissues, followed by RT-PCR for the viral NS5 gene. The viral RNA copy number was calculated from the standard curve plotted separately using in-vitro synthesised viral RNA from infectious cDNA clone. (G) - Predicted mRNA binding targets of HuR proteins, which may help in pathogenesis. (H) - IL-8 and TNF-alpha levels in mice liver tissues. Total RNA was isolated from infected mice liver tissues, followed by RT-PCR for the viral IL-8 and TNF-alpha levels and GAPDH RNA was taken as an internal control. (I) - IL-8 and TNF-alpha levels in HuR KO cells post virus infection. Total RNA was isolated from infected HEK293T and HEK293T HuR KO cells, followed by RT-PCR for the viral IL-8 and TNF-alpha levels, and GAPDH RNA was taken as an internal control. (J, K) - Western for HuR immunoprecipitation in mouse liver tissues at Day 3 post-infection. Immunoprecipitation was done with anti-HuR antibody. (L, M) - RT-PCR of IL-8 and TNF-alpha from the HuR immunoprecipitated fraction at Day 3 post-infection. Real-time PCR was done from the immunoprecipitated fraction and normalization was done with the Ct values of input RNA in each mice tissue. Students t-test was used for statistical analysis. P<0.05 = *, P<0.01 = **

HuR is a known RBP and an essential immunomodulatory protein that interacts and stabilizes many genes that regulate host immune responses, such as chemokines and cytokines. HuR is also known to shuttle from the nucleus to the cytoplasm via post-translational modifications, predominantly phosphorylation, under certain stress conditions such as oxidative stress and virus infection [15]. Upon DENV infection, the relocalization or phosphorylation of HuR protein was not detected, which is consistent with the previous observations of no relocalization for other vital factors such as PTB and DDX6 in Huh 7 cells [30–32]. This may be due to the cytoplasmic amount of the protein being sufficient to regulate the viral life cycle, as shown by the co-localization of HuR protein with the viral NS5 protein using digitonin as a cytoplasmic-specific permeabilization agent, as well as the silencing and overexpression results.

The interaction of HuR protein with the 5’UTR has predominantly been reported to play a role in viral RNA translation, as seen in the HCV virus [33]. In contrast, its interaction with the 3’UTR has a predominant role in replication, such as in the case of CVB3 [19]. The 3’UTR-specific interaction of HuR and the lack of any effect on the viral translation using the replicon system show that, in the context of DENV, it has a predominant role in viral RNA replication compared to cap-dependent translation. Interestingly, the predicted binding site of HuR overlaps with another known RBP, PTB, previously known to regulate viral RNA replication positively. This is evident in our study using gene-specific KO CRISPR cell lines of HuR and PTB. The overlapping interaction of HuR and PTB, along with the in vitro and ex vivo experiments, demonstrated that during infection i.e., from 24 h to 48 h when the viral RNA starts replicating, the displacement of HuR protein by PTB from the 3’UTR of the viral RNA due to higher affinity of PTB act as a molecular switch that helps the viral RNA peak post 24 h of infection.

DENV has previously been shown to exhibit both cap-dependent and cap-independent mechanisms of translation [7, 34, 35]. However, it has not been shown at which phase of the viral life cycle the viral RNA undergoes a transition from cap-dependent to cap-independent translation and which host factors are required in this process. Using reporter constructs, we were able to show that the cap-independent mode of the translation may be prominent at a later phase of the viral life cycle, i.e., 48 h post-infection, which can be attributed to a reduction in the levels of canonical translation factors[20] due to an overall reduction in global translation upon virus infection as shown by polysome analysis. The results indicated a specific role of 3’UTR in cap-independent translation, which corroborates the previous observations [7, 36]. Luciferase assays performed in the presence of virus infection and the cap inhibitor LY294002 in HuR KO cells show the role of HuR protein in regulating cap-independent translation, which may be due to the 3′UTR-binding activity of HuR protein that inhibits replication, thereby allowing more viral RNA to be available for translation, or possibly due to the dysregulation of host factors that are themselves regulated by HuR protein [37].

The infection in the AG129 mouse model also corroborated our cell culture data, showing higher levels of HuR protein when virus titers were low. In contrast, high virus titers led to downregulation of HuR protein levels, facilitating greater viral RNA replication. The mRNA targets of HuR protein, which have previously been shown to play important roles in the cytokine pathway [21], were also found to be dysregulated upon virus infection, inversely correlating with changes in HuR protein levels in both cell culture and mouse models. This suggests a destabilizing role for HuR in the regulation of cytokines upon DENV infection. Overall, this study uncovers how the virus cleverly uses host cell machinery at different stages of infection, demonstrating for the first time all the host factors present in the viral replication complex and elucidating the double role of an important RBP, HuR, in the DENV life cycle and pathogenesis, highlighting its potential as a target for antiviral therapies.

## Acknowledgments

We would like to acknowledge Prof. Ralf Bartenschlanger for sharing with us Dengue Su-genomic replicon (pFK SgDV-R2A). We would also like to acknowledge IISc DBT partnership and Mass spectrometry facility at NIBMG, Kalyani.

## Materials and Methods

### Isolation and Characterization of DRMs

To characterize the DRMs, membrane flotation assay was performed as described [38]. The cells were lysed in hypotonic buffer by passage 20 times through a 25-gauge needle. Centrifugation was used to remove cell debris and nuclei for five minutes at 1,000 × g. Cell lysates were mixed with 1.5 ml of 72% sucrose in low-salt buffer (LSB). The mixture was overlaid with 55% sucrose in LSB followed by 0.75 ml of 10% sucrose in LSB. The sucrose gradients were centrifuged at 38,000 rpm for 14 h at 4°C using a Beckman SW55 Ti rotor. Fractions of 500 μl were collected, and 100 μl from each fraction was resolved by SDS-PAGE followed by Western blotting. To analyze the proportion of proteins in the detergent-resistant membrane (DRM) fractions, cell lysates were treated with 1% NP-40 for 30 min on ice before ultracentrifugation.

### LC-MS/MS analysis

Protein extraction and trypsin digestion. A total of 100 μg protein was reconstituted in 50 mM ammonium bicarbonate buffer, reduced with 10 mM dithiothreitol for 1 hour at 65°C followed by alkylation with 40 mM iodoacetamide for 30 minutes at 37°C in dark. The proteome was hydrolysed overnight with MS-grade trypsin with final protease to protein ratio of 1:50 at 37 °C. Further the digested peptides were cleaned by solid phase extraction by using Sep-Pak Vac 1cc (50mg) C18 Cartridges (Waters Corporation). The peptide concentration was determined by BCA assay before being loaded onto analytical column.

### LC-MS/MS acquisition and database search

The proteomic dataset was acquired by using Ultimate3000 RSLC nano system online coupled to a QExactive Plus Orbitrap MS with an EASY nano-Spray interface (Thermo Fisher Scientific). The peptides were resolved on a C18 analytical column (2 μm, 100 Å particles, 75 μm × 50 cm), mobile phases for peptide separation (A, 0.1% FA in 5% acetonitrile; B, 0.1% FA in 100% acetonitrile) delivered by the RSLC nano system with flow rate 0.3 μL/min. The gradient time was 117 min, reaching 38% of B. The peptides were ionized in positive mode and acquired in full scan using top 15 method for precursor selection with a resolution 140,000, custom AGC target followed by 15 MS/MS scans with a resolution of 17,500, custom AGC target, mass isolation window of 1.4 m/z, and normalized collision energy at 29 eV. The proteomic data set was processed and analysed by using SEQUEST HT search algorithm in Proteome Discoverer v2.4 (Thermo Fisher Scientific) with a UniProt human database for protein identification and quantification across all samples. The search parameters included were oxidation of methionine (15.99 Dalton), and fixed modification of cysteine carbamidomethylation (57.021464 Dalton). Peptide identification was performed using a 10-ppm precursor ion tolerance and 0.02 Da for-product ions, peptide spectrum matches were adjusted to 1% false discovery rate. Proteins with at least 1 unique peptide were considered for further analysis.

### Cell lines and transfections

Huh7 cells were provided by the laboratory of Charles M. Rice, Rockefeller University, and Apath, LLC (New York, NY, USA) and maintained in Dulbecco’s modified Eagle’s medium (DMEM; Sigma) with 10% fetal bovine serum (Gibco, Invitrogen). Transfection was carried out using Lipofectamine 2000 (Invitrogen) in accordance with the manufacturer’s protocol. For cell culture experiments, Huh7 cells were infected with *Dengue* serotype 2 at MOI-1. Uninfected Huh7 cells were used as the control.

### Dengue infection on Huh7 cells

Huh7 cells (70% confluent monolayer) were infected with MOI-1 of the dengue serotype 2 virus in serum-depleted DMEM Two hours after infection, DMEM supplemented with 10% fetal bovine serum was added. Cells were harvested after 24 h, 48 h, and 72 h using TRI reagent (Sigma) for total RNA isolation and qRT-PCR.

### Quantitative real-time polymerase chain reaction (qRT-PCR)

SYBR Green mRNA Assay System was used for Real Time PCR. The RT reaction was carried out in three stages. The first phase included 600 ng of total RNA, 0.5 µl of RNase Inhibitor, 2 µl of mRNA-specific reverse primer (10 mM), and 2 µl of GAPDH reverse primer (10 mM). Snap cooling in ice, 75 °C, and 5 minutes were the reaction conditions. This reaction mixture was then supplemented with 0.1 µl of reverse transcriptase enzyme, 2 µl of 10× RT buffer, and 2 µl of 10 mM dNTPs mix. Five minutes at 25 °C, one hour at 42 °C, and ten minutes at 75 °C were the final reaction conditions. After that, RT-PCR was carried out in a total reaction volume of 10 µl containing 5 µl of 2× Master Mix, 1 µl of mRNA-specific forward primer (10 mM), 1 µl of mRNA-specific reverse primer (10 mM), 0.2 µl of Rox, and 2 µl of RT product. The reaction conditions were 95 °C for 10 min, followed by 40 cycles of 95 °C for 30 s, 61 °C for 30 s, and 72 °C for 30 s.

### siRNA transfection

Briefly, 14 h after seeding of cells, 150 nM siRNA targeting HuR, and a nontargeting siRNA (Dharmacon) was transfected using Lipofectamine 2000 transfection reagent in Opti-MEM (Invitrogen). At 96 h posttransfection the cells were harvested, and the extracts were used for Western blot analysis as described below. In the case of transient transfections, siRNA was transfected first, and Dengue virus was infected 16 h later. Cells were harvested at the time points after Dengue infection as indicated on the figures.

### Immunofluorescence staining

For immunofluorescence staining, ∼0.06 × 10^6^ Huh 7 cells were seeded in a 24-well plate on coverslips for 14 h, followed by infection with Dengue virus (MOI=1). At 48 h post-infection, cells were washed twice with 1× phosphate-buffered saline (PBS) and fixed using 4% PFA at room temperature (RT) for 10min. Permeabilization was done with 0.1% Triton X-100 or 10ug/ml digitonin for 10 min at room temperature, after one hour of incubation with 3% bovine serum albumin (BSA) at 37°C, the cells were treated with the antibodies shown in the figure for two hours at 4°C. Alexa-488-conjugated anti-mouse or Alexa-647 conjugated anti-rabbit secondary antibodies were then used for 30 minutes (Invitrogen) to identify the cells. A Zeiss microscope was used to take the pictures, and the Zeiss LSM or ZEN software program was used to analyze the images. The overlap coefficient was used to measure colocalization; a value of 0 denotes no colocalization, while a value of 1 denotes full colocalization. For each experiment, over 60 cells were examined.

### Western blot analysis

Protein concentration was measured using the Bradford test (Bio-Rad), and equal volumes of cell extracts were separated using SDS–12% PAGE and then put onto a nitrocellulose membrane (Pall Corporation).Using the appropriate secondary antibody (horseradish peroxidase-conjugated anti-mouse or anti-rabbit IgG [Sigma]), the samples were analyzed by Western blotting using the desired antibodies, which included anti-HuR antibody (3A2; Santa Cruz), anti-NS5B antibody (GTX103350; Genetex), anti-La antibody (ab75927; Abcam), or anti-PTB antibody (Calbiochem). To ensure that entire cell extracts loaded equally, a mouse monoclonal anti-β-actin peroxidase-conjugated antibody (A3854; Sigma) was employed as a control. An Immobilon Western system (Millipore) was used to detect antibody complexes.

### *In vitro* transcription

Runoff transcription processes were used to in vitro transcribe RNAs from various linearized plasmid constructs using T7 promoters. Xba1 was used to linearize pcDNA3 vectors carrying the HCV 5′ UTR or HCV 3′ UTR. After being electrophoresed on agarose gels and extracted using a Qiagen gel elution kit, the linear DNA samples were utilized as templates for the production of unlabeled or 32P-labeled RNA utilizing [32P] UTP (PerkinElmer Life Sciences) and T7 RNA polymerase (Fermentas). 2.5 μg of linear template DNA was used in the transcription reaction, which was conducted for 1.5 hours at 37°C using normal conditions (Fermentas protocol). The RNA was resuspended in 20 μl of nuclease-free water following alcohol precipitation. One microliter of the radiolabeled RNA sample was spotted onto DE81 filter paper, washed with phosphate buffer, and dried, and the incorporated radioactivity was measured using a liquid scintillation counter.

### Protein purification

In Escherichia coli BL21(DE3) cells that had been transformed using the proper pET28a vectors, recombinant HuR, PTB, and La proteins were produced. At an optical density of 0.6 at 660 nm, 0.5 mM isopropyl-1-thio-β-d-galactopyranoside (IPTG) was used to promote the expression of recombinant HuR, PTB, and La. The cells were then allowed to proliferate for an additional five hours. Sonication was used to break up the cells on ice after they had been pelleted and resuspended in lysis buffer (50 mM Tris, pH 7.5, 300 mM NaCl, 0.1 mM phenylmethylsulfonyl fluoride [PMSF]). Every step that followed was done at 4°C. After centrifuging the lysates for 30 minutes at 10,000 rpm, they were incubated for 6 hours with rocking in a Ni-nitrilotriacetic acid (NTA)-agarose slurry (Qiagen). The flowthrough was disposed of, and the lysate was put onto a column. 20 ml of wash buffer (50 mM Tris, pH 7.5, 300 mM NaCl, and 40 mM imidazole) was used to wash the column. 500 μl of an elution solution containing 500 mM imidazole was used to elute the bound protein. Following a 4- to 6-hour dialyzation in ten times the volume of dialysis buffer (50 mM Tris, pH 7.4, 100 mM KCl, 7 mM β-mercaptoethanol [β-ME], 20% glycerol), the eluted proteins were aliquoted and kept in a freezer at -70°C.

### Preparation of S10 extracts

The preparation of S10 extracts followed as described by Ray et al (16). In summary, 10% FBS (Gibco, Invitrogen) was added to DMEM (Sigma) to support the growth of Huh7 or replicon cells. Following three rounds of washing with cold isotonic buffer (35 mM HEPES, pH 7.4, 146 mM NaCl, and 11 mM glucose), a monolayer of cells was harvested, pelleted down, and resuspended in 1.5× packed cell volume of hypotonic buffer (10 mM HEPES, pH 7.4, 15 mM KCl, 1.5 mM Mg-acetate, and 6 mM β-ME). The cells were then allowed to swell for 10 minutes on ice. After that, the cells were sent to a Down’s homogenizer and given 50 ice strokes to break them up. 20 mM HEPES, 1.2 M KCl, 50 mM Mg-acetate, and 60 mM β-mercaptoethanol comprised the 1× incubation buffer in which the lysate was incubated for 10 minutes. To obtain the cytoplasmic extract (S10 supernatant), the lysate was centrifuged at 10,000 × g for 30 minutes at 4°C. One hundred liters of dialysis buffer (10 mM HEPES, 90 mM KCl, 1.5 mM Mg-acetate, 7 mM β-ME, and 20% glycerol) were dialyzed against the supernatant for 2 to 4 hours.

### UV-induced cross-linking of proteins with RNA and immunoprecipitation (IP) assays

The procedure outlined by Ray and Das (16) for UV-induced cross-linking was used. Briefly, in 1× RNA binding buffer (5 mM HEPES, pH 7.6, 25 mM KCl, 2 mM MgCl2, 3.8% glycerol, 2 mM dithiothreitol [DTT], and 0.1 mM EDTA), α-32P-labeled RNA probes were allowed to form complexes with S10 extracts or with recombinant proteins before being exposed to UV light for 20 minutes. The mixture was separated on an SDS–10% polyacrylamide gel (SDS-PAGE), treated with 30 μg of RNase A (Sigma), and then subjected to phosphorimaging analysis. RNase A-treated reaction mixtures (30 μg of total protein) were prepared for immunoprecipitation (IP) using polysome lysis buffer (100 mM KCl, 5 mM MgCl2, 10 mM HEPES, pH 7.0, 0.5% NP-40, 1 mM DTT, and 100 U/ml RNasin) up to 500 μl. with polysome lysis buffer (100 mM KCl, 5 mM MgCl_2_, 10 mM HEPES, pH 7.0, 0.5% NP-40, 1 mM DTT, 100 U/ml RNasin) and precleared with protein G-Sepharose beads for 1 h at 4°C. To pellet the beads, the samples were spun at 1,000 × g for two minutes, and the supernatant was then discarded. Protein G-Sepharose beads were added to the precleared lysates and incubated for three hours at 4°C with continuous mixing using a rotator device. They were incubated with 2 μg of anti-HuR antibody (Santa Cruz) for an entire night at 4°C in a total volume of 200 μl of polysome lysis buffer. Polysome lysis buffer was used to wash the beads four times. After the immunoprecipitated protein was released from the beads by boiling them in SDS sample buffer, the supernatant was electrophoresed on an SDS–10% PAGE gel. Autoradiography was used to create, expose, and dry the gel.

### IP-RT assay

An experiment known as ribonucleoprotein complex immunoprecipitation (RNP IP) was used to evaluate the relationship between HuR and PTB and Dengue RNA. In short, IP was carried out using 2 μg of either anti-HuR and anti-PTB antibody or IgG isotype control antibody (Imgenex) after whole-cell lysates were produced in polysome lysis buffer. Before being treated with the corresponding antibodies in polysome lysis buffer for 16 hours at 4°C, Protein G beads (Sigma) were preblocked with 0.4% BSA for 30 minutes at room temperature. For each reaction volume, one milligram of cell lysate was employed. After lysates were precleared with protein G beads for one hour at 4°C, they were treated for four hours at 4°C with the target antibody. After four rounds of washing with polysome lysis buffer, the beads were treated with 0.1% SDS and 30 μg of proteinase K at 50°C for 30 min, followed by RNA isolation and RT-PCR to detect the presence of Dengue RNA. The ct values of the input RNA from each condition was taken as a normalization control.

### Sucrose gradient Polysome fractionation assay

To identify the association of viral RNA in polysomal fraction and effect of virus infection on global translation, sucrose gradient polysome fractionation was performed. After 48h post infection, cells were treated with 100 µg/ml cycloheximide for 15min at 37°C. Followed by that cells were washed with PBS and thereafter wash with 1x hypotonic buffer (5 mM MgCl2, 5 mM TRIS-HCl pH-7.5, and 1.5 mM KCl). 100 µg/ml cycloheximide was added for all buffers used for sample processing. Cells were scraped in lysis buffer (5 mM TRIS464 HCl pH-7.5, 5 mM MgCl2, 1.5 mM KCl, 100 µg/ml cycloheximide, 1mM DTT, 200 U/ml 465 RNase in from Promega, 0.5% Sodium-deoxycholate, 0.5% Triton X -100, 200µg t-RNA and 466 1X protease inhibitor cocktail) and incubated at ice for 20mins. After completion of lysis, the KCL was added in the lysate to adjust KCL concentration to 150mM and followed by that supernatant was collected after spinning the lysate for 8min at 3000 g at 4°C. 10-50% w/v sucrose gradient was prepared and supernatant was added from the top and samples were ultracentrifuged for 2h at 36000rpm at 4°C in sw41 rotor (Backman). Density gradient fractionation system was further used to fractionate the gradients at a flow rate of 0.3mm/sec in different microfuge tubes. UV detector generates polysome profiles, which further used to distinguish the tubes containing 40s,60s, monosomes and polysomes.

### Luciferase assays

Transfected cells were isolated and lysed with Promega’s 1× passive lysis buffer. The cell lysates underwent a 10-minute, 10,000-rpm centrifugation at 4°C. After collecting the supernatant, the Dual Luciferase kit (Promega) was used to take the luciferase reading in accordance with the manufacturer’s instructions. The measurements were adjusted for total protein for sub-genomic reporter and by R luc for 5’Luc and 5’Luc3’ reporter constructs.

### AG129 mice infection

All animal experiments were approved by BRIC-Rajiv Gandhi Centre for Biotechnology (BRIC-RGCB) Institutional Animal Ethics Committee (IAEC) (Approval No. (IAEC/911/ SREE/2022). AG129 mice (IFN-α/β and IFN-γ receptor knockouts) were procured from B&K Universal (UK), bred and maintained under specific-pathogen-free conditions.Three- to four-week-old AG129 mice were infected subcutaneously with 1 × 10⁴ plaque-forming units (PFU) of DENV-2 (strain RGCB880 35P) in a total volume of 500 μl. Mock-infected control animals were administered an equivalent volume of heat-inactivated virus as described [39] . Post-infection, mice were daily monitored for clinical symptoms including ruffled fur, hunched back, lethargy, edema with closed eyes and weight loss. Whole blood was collected by cardiac puncture under isoflurane anesthesia, on day 3 post-infection (early symptomatic phase) and day 6 post-infection (peak symptomatic phase) from both infected and mock-infected control animals. Blood samples were further processed for serum isolation. Immediately following blood collection, mice were sacrificed, and tissues including liver, spleen, and brain were harvested and preserved in RNA later for subsequent molecular analysis.

## Supporting information

**S1 Table.**

**S2 Table.**

**S1 Fig.**
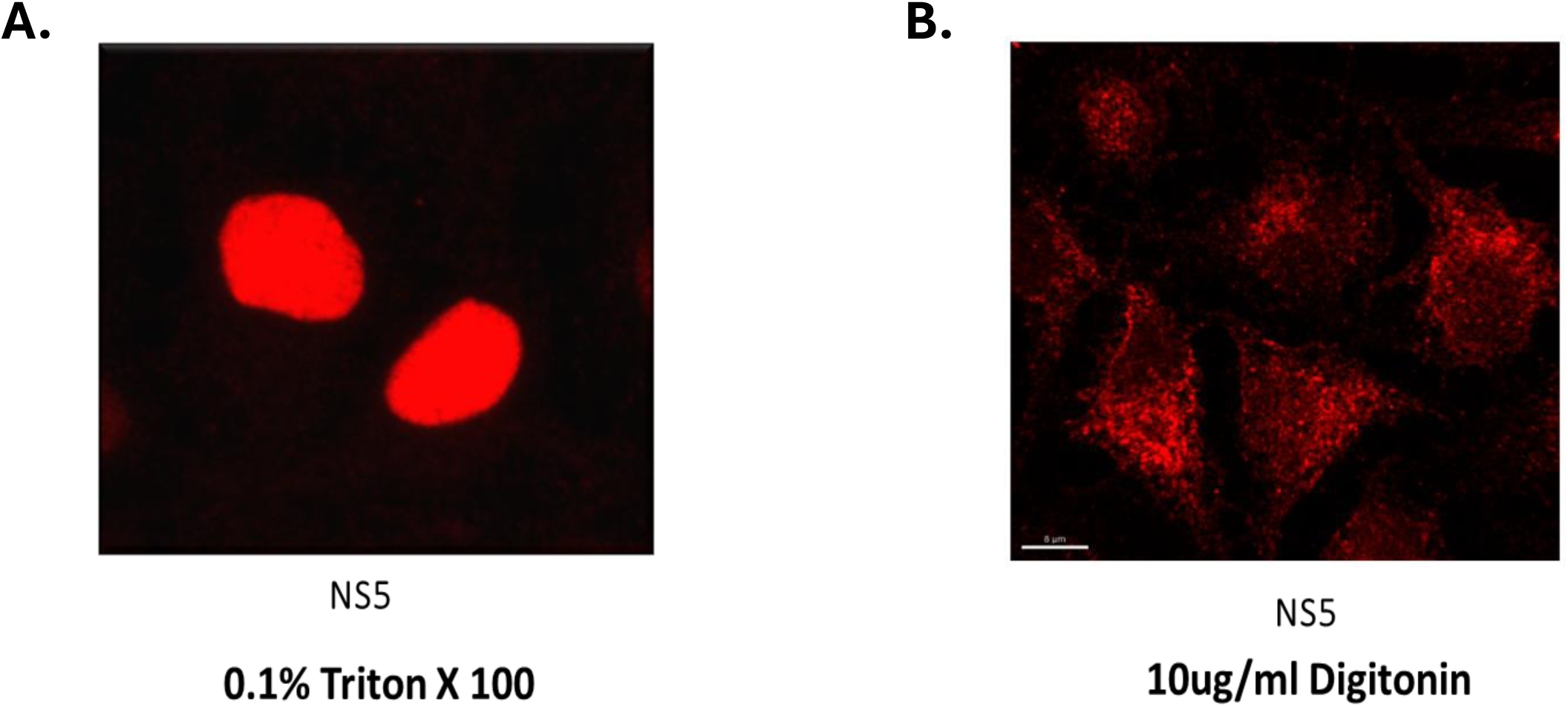
Standardization of Nuc-cyto levels of a single protein using different permeabilization.

**S2 Fig.**
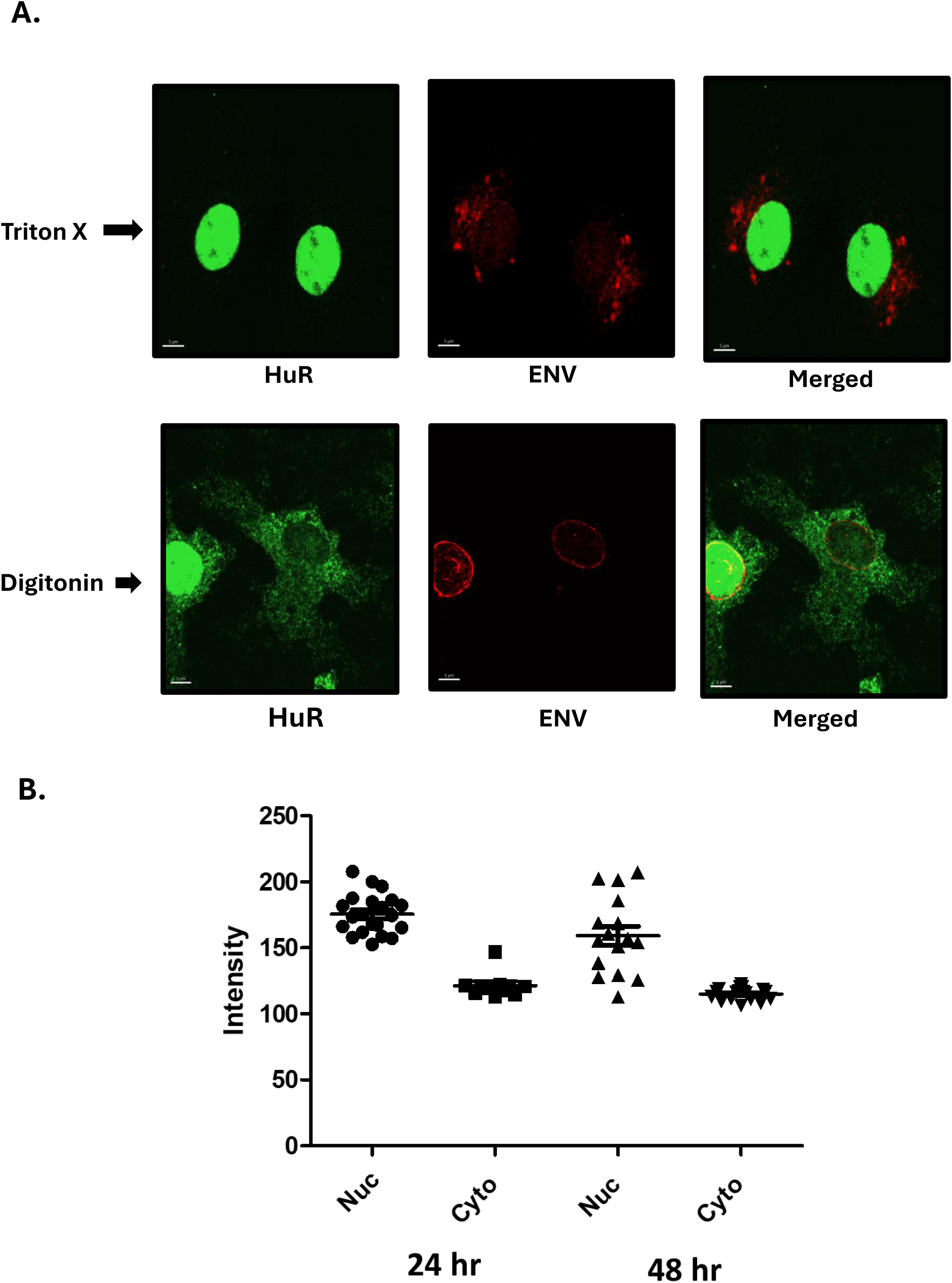
(A, B) **–** Nuc-Cyto levels of HuR protein at a single infected cell resolution using digitonin-mediated permeabilization

**S3 Fig.**
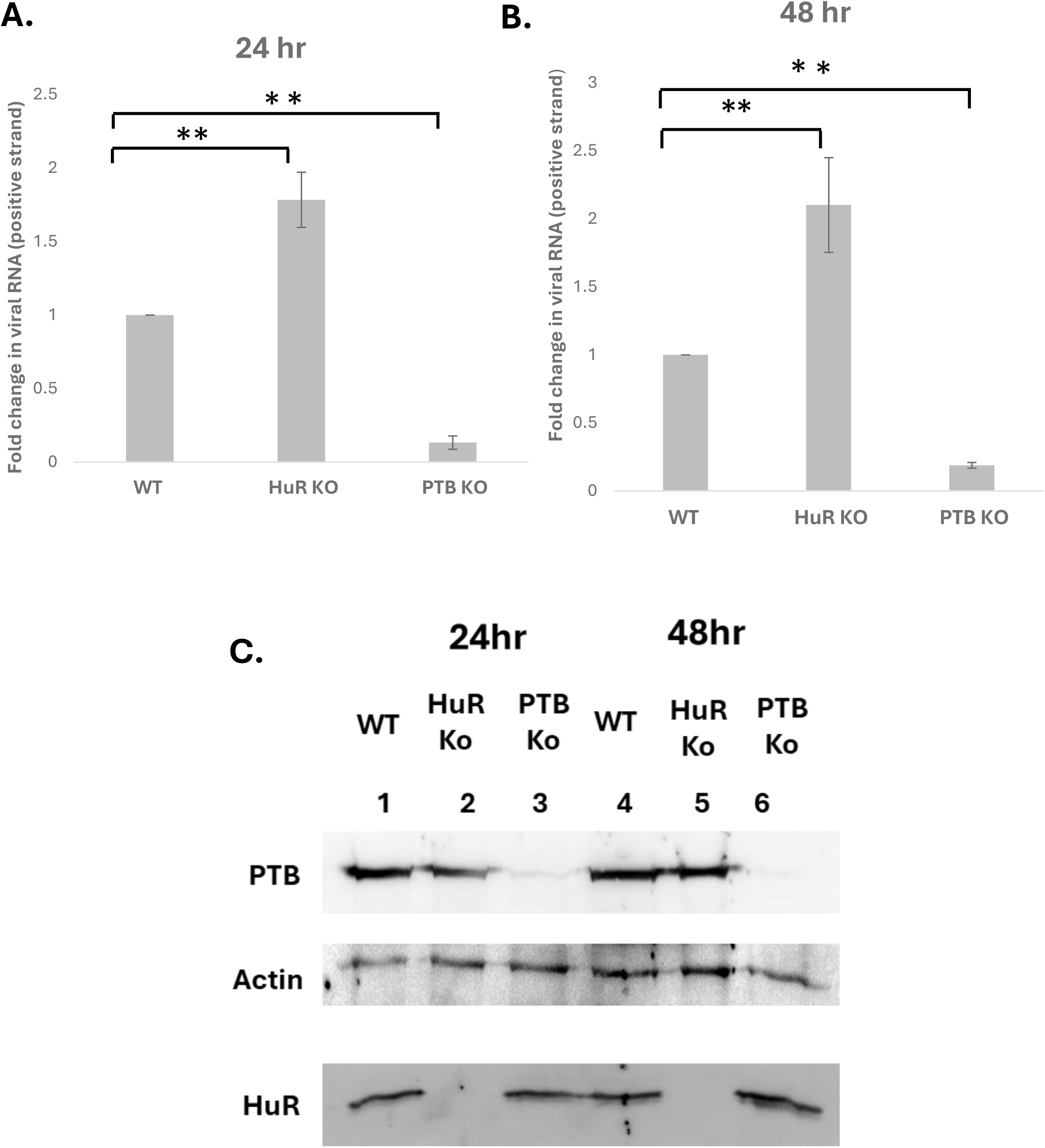
(A, B) – Fold change in virus levels in HuR and PTB KO cells. (C) Validation of KO cells using western blotting.

**S4 Fig.**
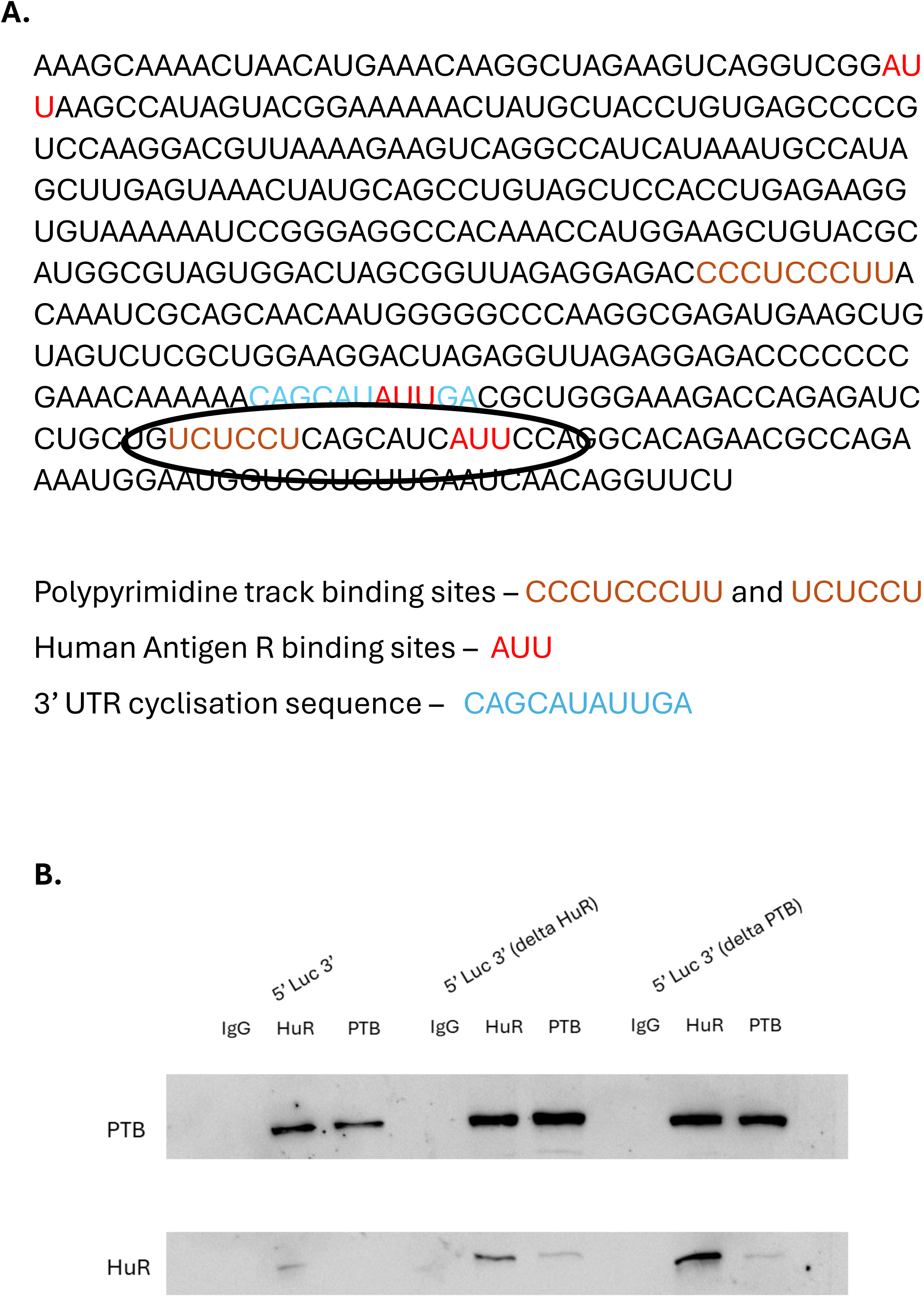
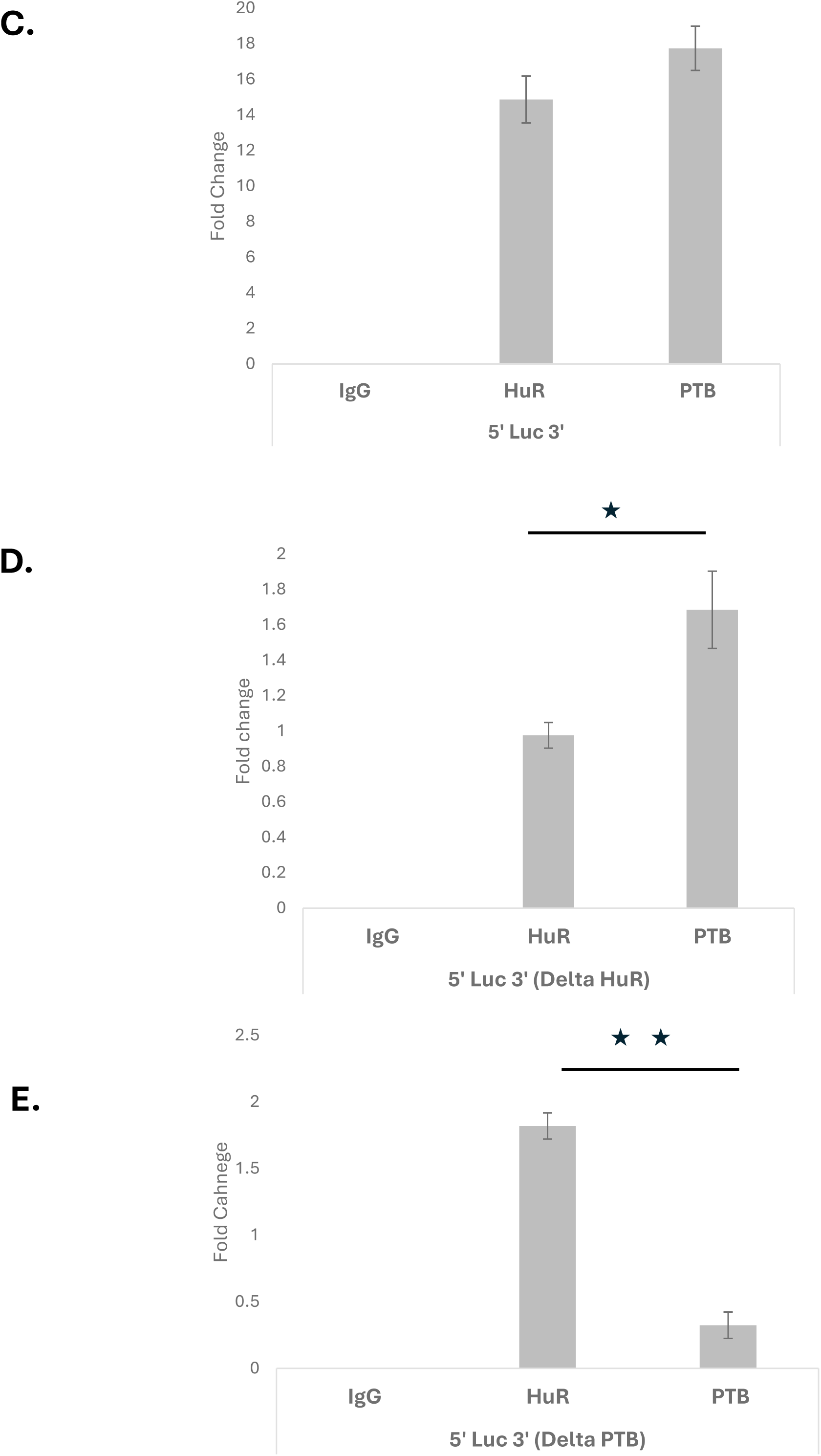
(A) – HuR and PTB binding site at the 3’UTR of viral RNA. (B) – Western blotting showing HuR and PTB IP post-transfection. (C-E) – RT-PCR from HuR and PTB immunoprecipitated RNA upon transfection of WT and delta HuR and delta PTB reporter constructs.

**S5 Fig.**
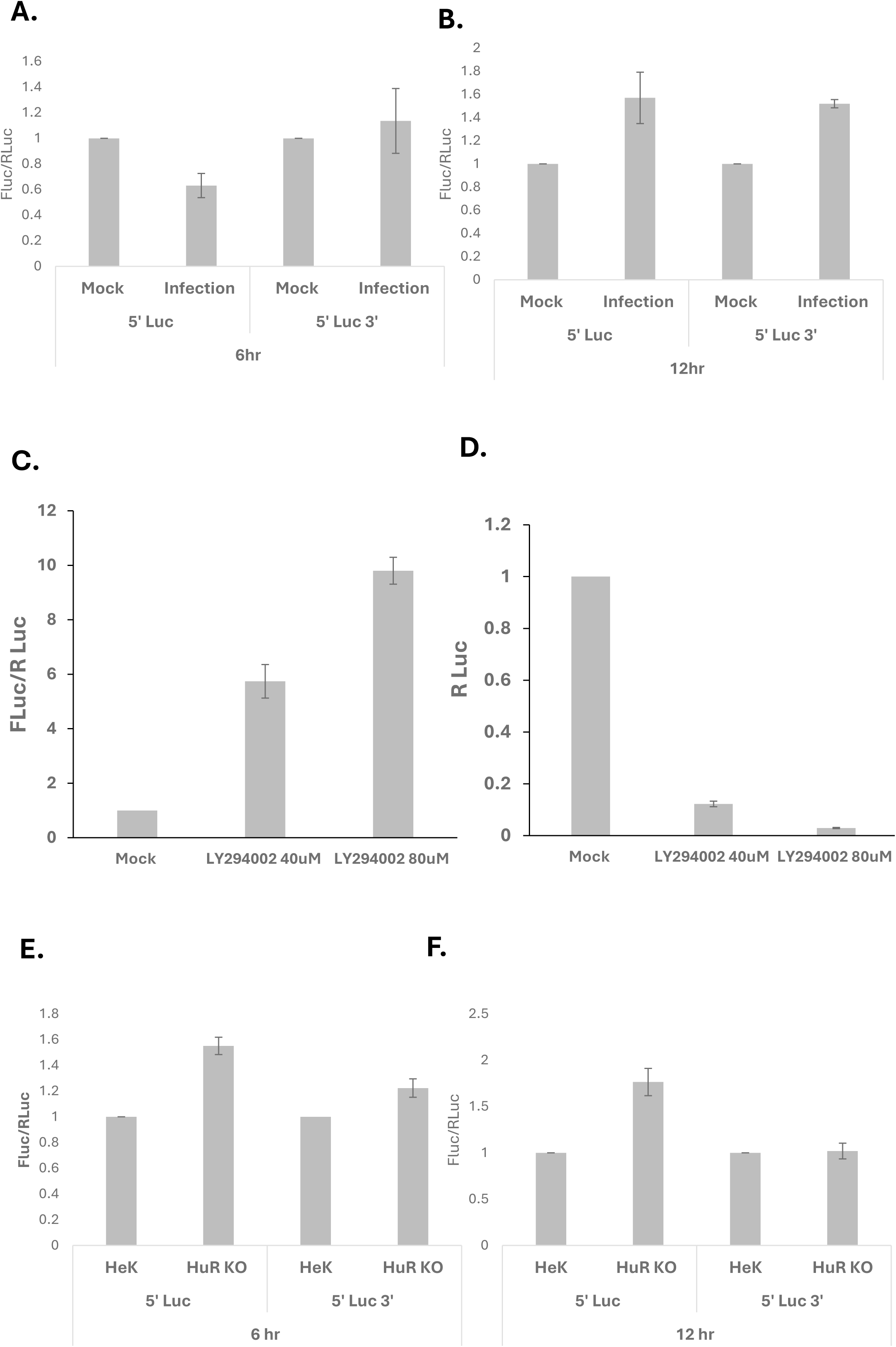
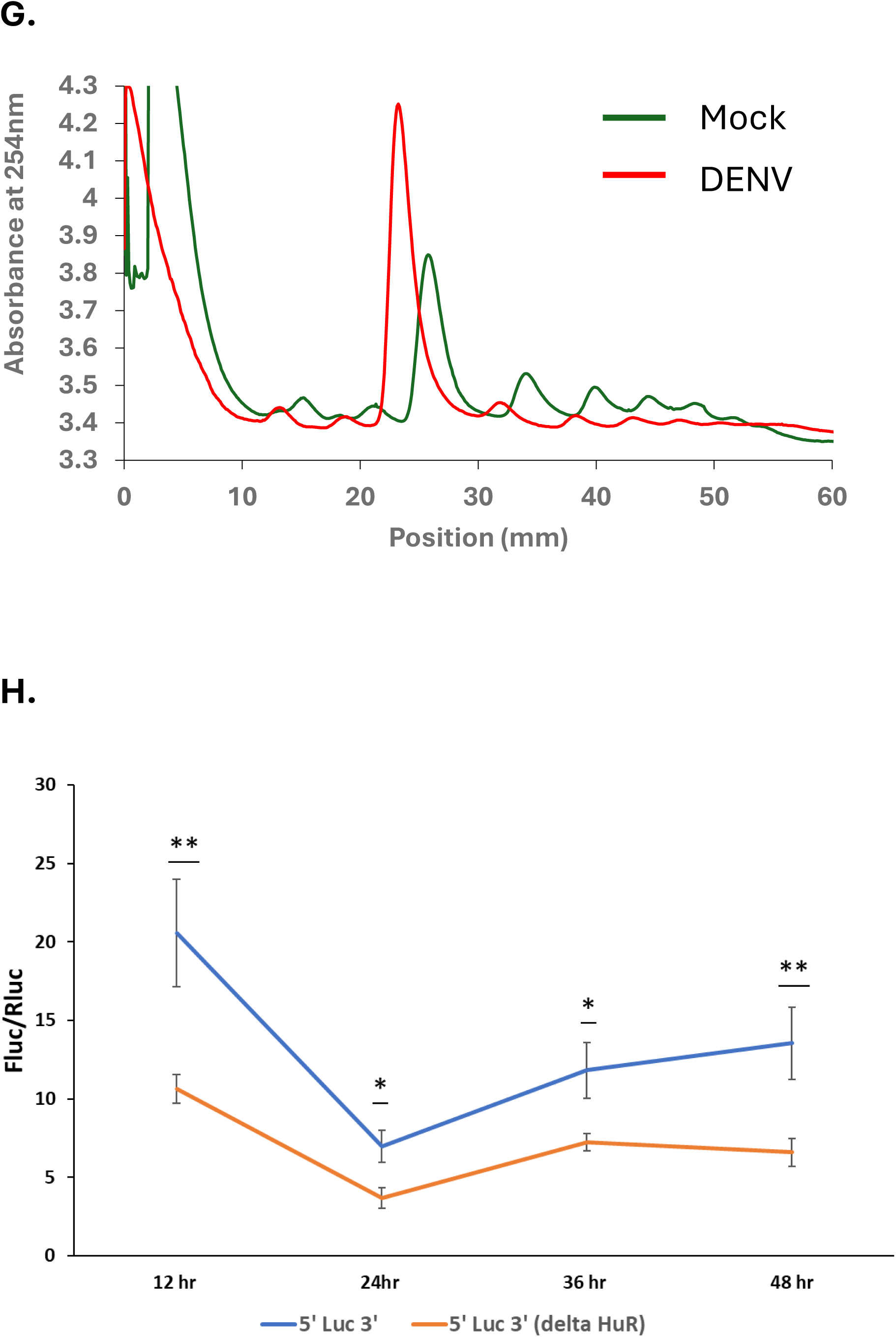
(A, B) – Relative Luc activity for 6hr and 12hr in the background of Dengue infection. (C, D) – Standardization of cap-independent translation in the presence of cap inhibitor LY294002. (E, F) – Relative Luc activity for 6hr and 12hr using cap inhibitor LY294002. (G) – Polysome profiling at 48hr post-infection in HuH 7 cells. (H) – Relative Luc activity of 5’Luc 3’ and 5’Luc3’(delta HuR) in the background of Dengue infection.

**S6 Fig.**
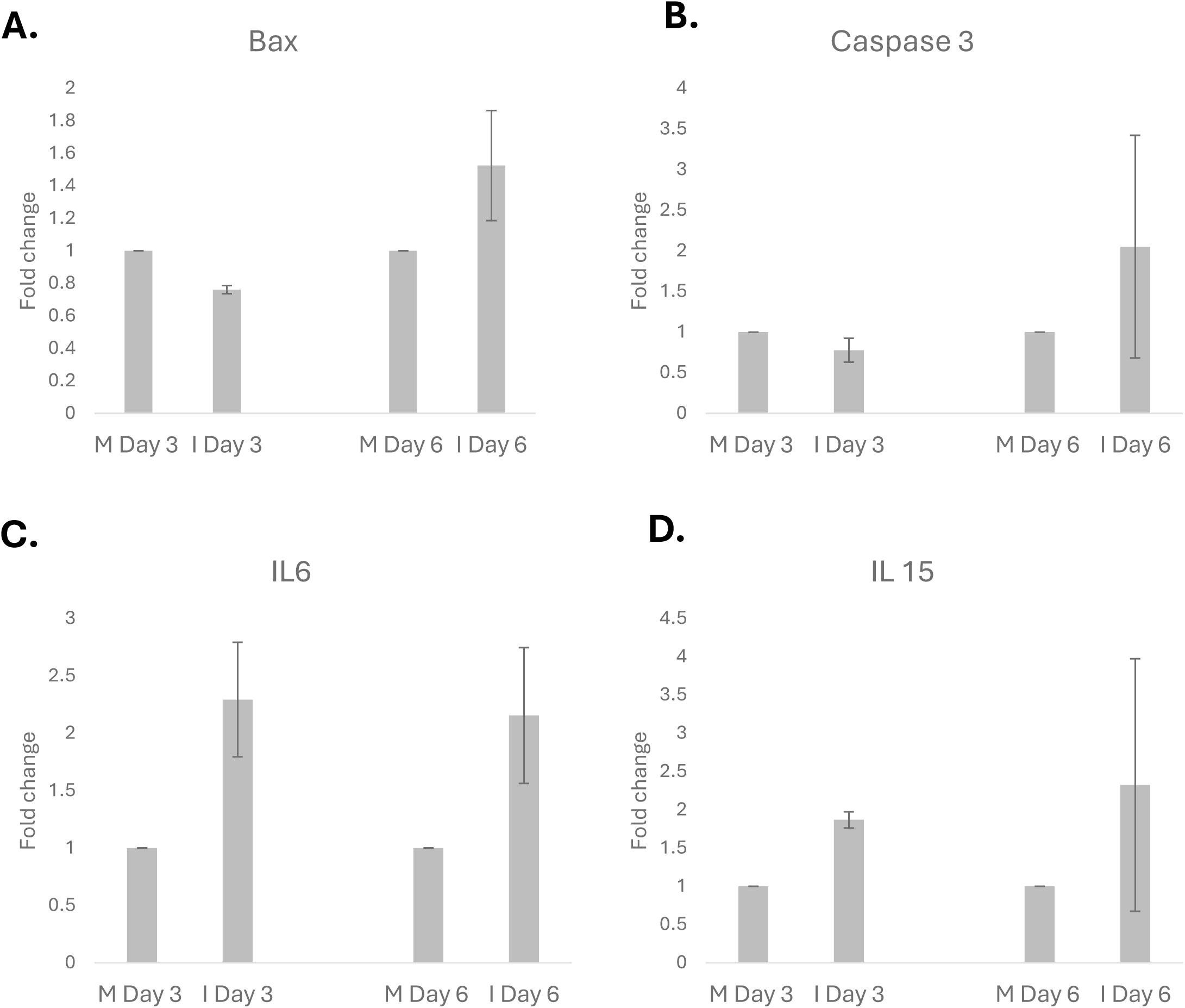
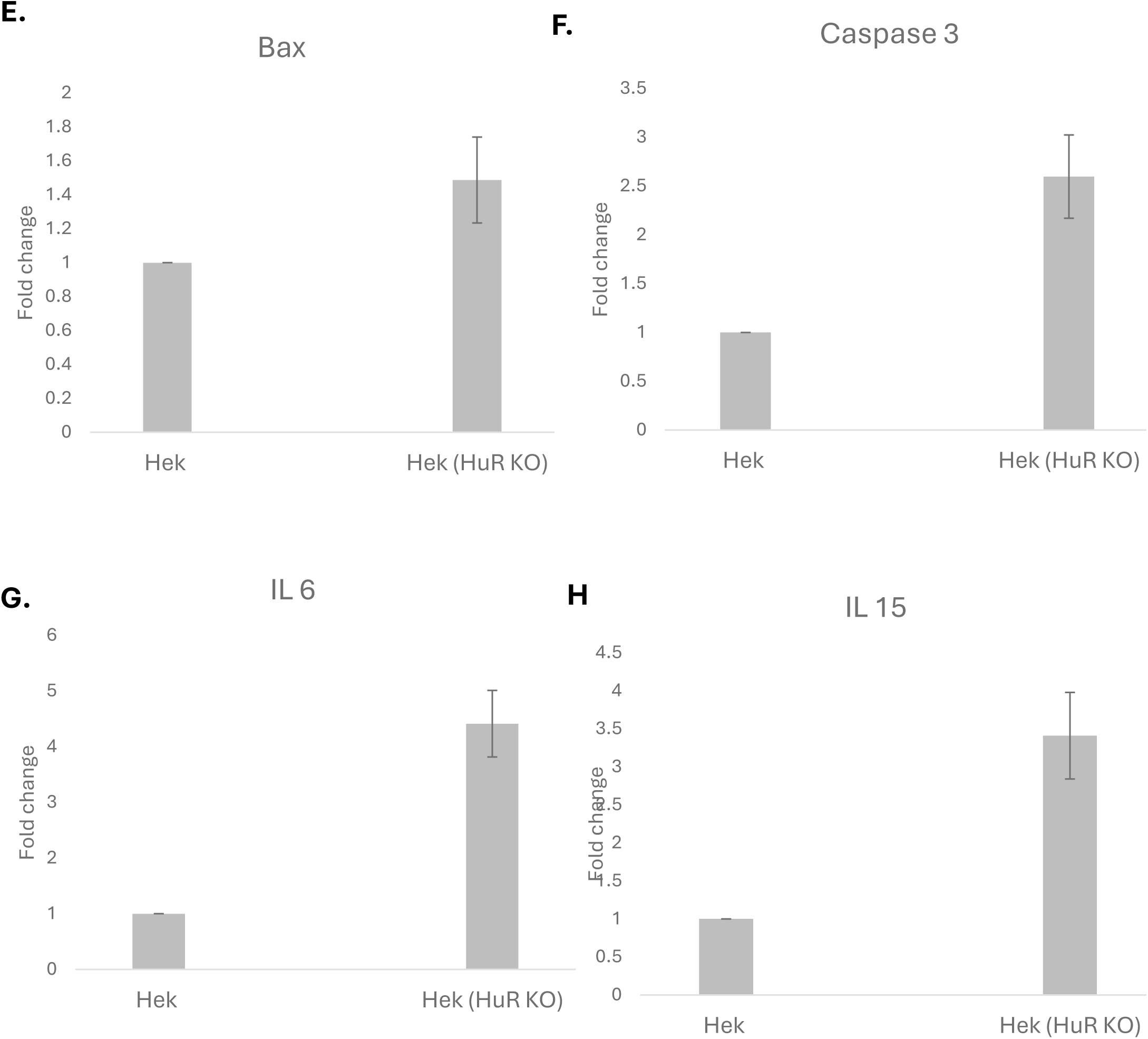
(A-D) - Validation of HuR targets in mice liver tissues. (E-H) - Validation of HuR binding mRNA targets in HuR KO cells.

